# Fibroblasts close a void in free space by a purse-string mechanism

**DOI:** 10.1101/2020.08.15.250811

**Authors:** Avelino Dos Santos Da Costa, Ramesh Subbiah, Seung Ja Oh, Hyun Tae Jeong, Jung-Im Na, Kwideok Park, Jennifer Hyunjong Shin, In-Suk Choi

## Abstract

How stromal cells fill voids in wounded tissue remains one of the most fundamental questions in regenerative medicine. Fibroblasts are known to fill voids by depositing extracellular matrix (ECM) proteins while migrating towards the wound site; however, their ability to adopt an epithelial-like purse-string behaviour remains unexplored. Here, we fabricated an artificial wound with a deep void space to investigate fibroblasts’ behaviour during gap closure. We found that fibroblasts can form a free-standing bridge on deep microvoids and consequently close the void through the purse-string contraction, which was previously believed to be exclusively an epithelial wound closure mechanism. The results also revealed that the fibroblast gap closure in our fabricated 3D artificial wound depends on myosin II-mediated contractility and intercellular adherent junctions. Our study reveals that stromal cells can gain the structural features of epithelial cells, namely, intercellular contractile rings, to fulfil their functions under the specific microenvironmental conditions of tissue repair. Furthermore, fibroblasts can close artificial wounds with gap widths up to 300 μm, approximately twice as large as the critical epithelial gap closure size on non-adherent substrates. Fibroblasts exhibited a zip-up gap closure mechanism with a geometrical size effect. These findings reveal a new mechanism for gap closure by stromal cells during wound healing and pave a way to groundbreaking therapeutic strategies for tissue repair.

## Introduction

Gaps and holes in constantly remodelling living tissue are continually formed and closed in multicellular tissues during morphogenesis, tissue injury and repair, and the extrusion of apoptotic cells. In the event of injury, the open wound needs to be closed promptly to restore normal physiological tissue function^1,2^. Additionally, the cellular behaviour in the microscopic level of gap closure recently gained further attention from dermatologists, especially for skin resurfacing such as fractional laser treatments^3^. These treatments deliver thousands of microbeams into the skin, which intentionally create microscopic thermal wounds that the shape of wounds differs with laser source and parameters^4,5^. Consequently, geometrical size effects on stromal cells will help design optimal microscopic wounds for the desired healing process. In order to understand healing mechanisms at the cellular level, studies have employed two-dimensional (2D) wound models by scratching or patterning voids on planar substrates^6–10^. Two well-known distinct mechanisms have been actively studied, the active crawling of cells at the lesion periphery and purse-string-like constriction by supracellular actomyosin cables^7,11–13^. Depending on the nature and physicochemical environment of the wound, the relative contributions of these two distinct modes can vary. For epithelial wounds, both of these modes of closure play significant roles when an extracellular matrix (ECM) is present on the substrate. While smaller gaps are closed effectively by purse-string-like contraction, the closure of larger gaps utilizes cell crawling with active lamellipodia protrusion as the predominant mechanism. When the voids are non-adherent with no ECM for the cells to bind to, epithelial cells are not able to anchor to the substrate but are forced to form free-standing epithelial sheets adjoined by stable cell-cell adhesions over the non-adherent substrates^9,14,15^.

Unlike epithelial cells, mesenchymal cells, represented by fibroblasts, mainly employ active crawling to fill the voided region on 2D substrates in the presence of ECM^16–19^. Therefore, without the ability to engage in active crawling, as on a non-adherent substrate, the fibroblasts fail to close the wound by forming free-standing sheets like epithelial cells due to lack of stable intercellular junctions.^14^. Recent reports have suggested that fibroblasts may close void spaces via cytoskeletal contractility and interaction of cell-ECM adhesion^16^. Moreover, fibroblasts cultured in a 3D-engineered cleft utilize supracellular circumferential actin cables to generate centripetal tensions and close the cleft^20^. However, fibroblasts’ ability to adopt an epithelial-like purse-string behaviour to induce gap closure remains unexplored.

In this study, we developed a 3D microvoid system to study void closure mechanisms utilizing NIH 3T3 mouse fibroblasts. Our fabricated deep microvoid system has a thickness of 110 μm, simulating a free-standing gap. During the course of gap closure, no lamellopodia protrusions were observed from the cells advancing on the front line, indicating no involvement of crawling mechanisms. Unlike previous reports, stromal cells managed to sense the structural features of the microvoid and adopted a purse-string contraction for gap closure. Wound closure by purse-string contraction has been believed to be a *bona fide* mechanism of epithelial cells. However, this study clearly demonstrates that stromal cells, represented by fibroblasts, can reorganize their actin filaments along the periphery that span multiple cells to close the gap by purse-string contraction.

### Closure of 3D microvoid as artificial wound by stromal cells

Artificial wound-mimicking 3D microvoids were fabricated with different gap sizes (100, 200, and 300 μm) using a polydimethylsiloxane (PDMS) substrate (Supplementary Fig. 1**a** and **b**). Each microvoid had a quarter-circle at each corner, referred to as the curvature of the substrate corner (κ_corner_) with a radius of 50 μm and had a sharp-edge morphology with a thickness of 110 μm (Supplementary Fig. 1**c**). The substrate was coated with polydopamine (PDA) (Supplementary Fig. 2). Cells were seeded on the base of 3D microvoids as a reservoir of cells, and with the progression of time, they formed bridges that initiated at the κ_corner_ and naturally filled the voids in a free-standing manner (Fig. 1**a**). During the course of void closure, we did not observe the protrusion of lamellipodia from advancing cells, indicating the lack of a crawling mechanism in our system (Fig. 1**b, c**). Moreover, closure occurred not only on smaller gaps (100 μm) but also on larger gaps up to 300 μm (Supplementary Movies 1-3). Additionally, cells at the periphery of the advancing front were circumferentially aligned, as evidenced by F-actin and nuclei morphology (Fig. 1**c** and Supplementary Fig. 3**a**) compared with those on the flat base (Supplementary Fig. 3**c**). We found that the degree of cell alignment, marked by the nucleus alignment, was proportional to the tension generated during gap closure (Supplementary Fig. 3**b**); inversely, the tension was reduced when the gap was fully closed. During the gap closure, we observed that nuclei of advancing cells (region *a*) displayed a significantly higher aspect ratio (AR) than those in other regions (regions *b-d*) (Supplementary Fig. 4), supporting our notion that the front cells require more tension for the closure of gaps in the 3D microvoids. The void areas decreased linearly with time with average rates of 6.2 ± 1.03 μm/hr, 2.12 ± 0.38 μm/hr and 1.26 ± 0.42 μm/hr for initial gap sizes of 100 μm, 200 μm, and 300 μm, respectively. Regardless of the initial gap size, the rate of closure did not vary significantly (Fig. 1**d**). The SEM image evidently showed bridge formation that partially covered the microvoid (Fig. 1**e**). Taken together, these results indicate that stromal cells form a bridge at the corners of the microvoids and eventually close the voids by filling the gaps without any supporting matrix.

**Fig. 1.**
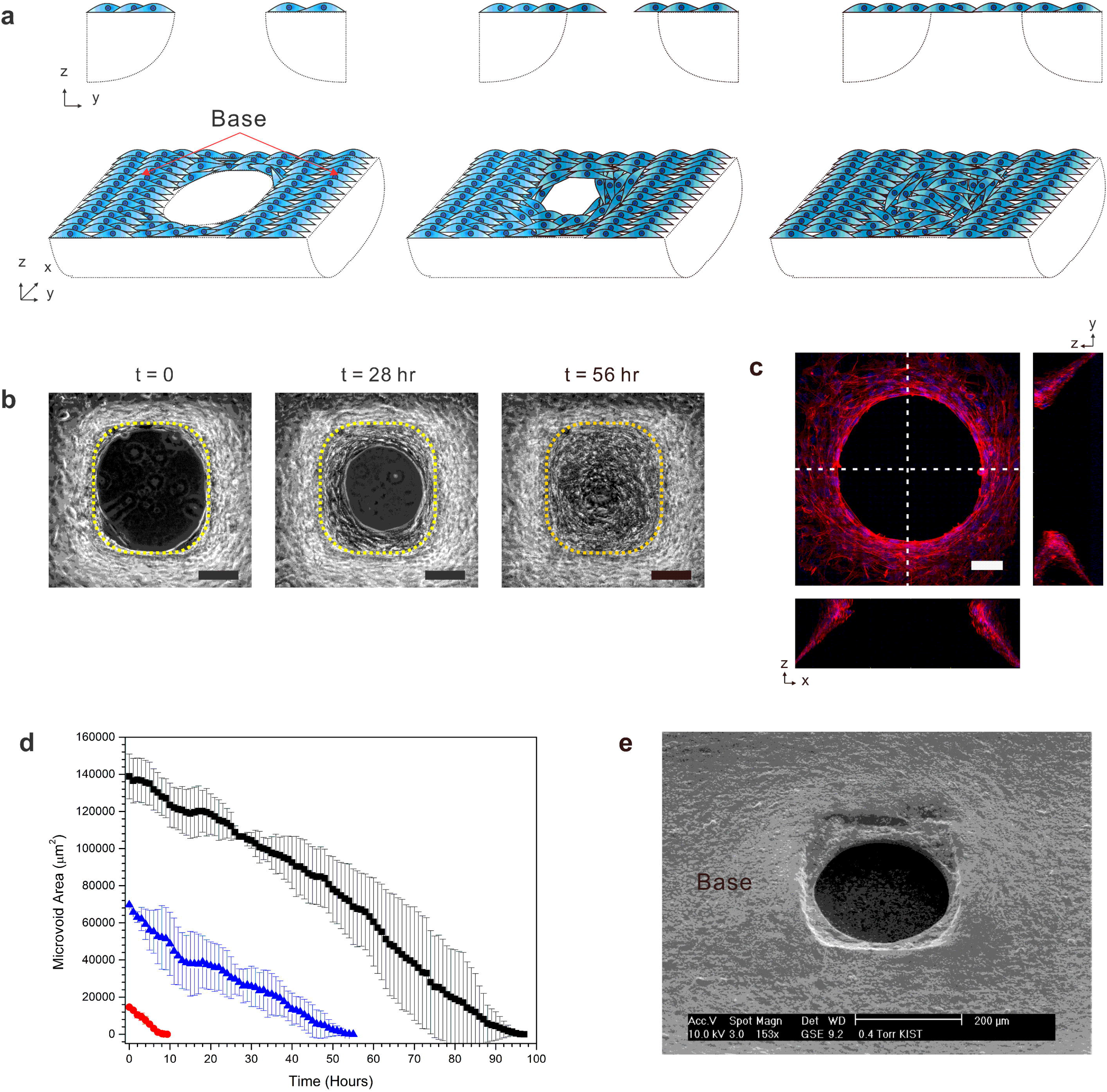
Fibroblasts close the void in 3D microvoids. a, Schematic illustrations of cross-sectional (*top row*) and tilted (*bottom row*) views of NIH 3T3 cells that formed a bridge and then closed a void over time. b, Phase contrast images of NIH 3T3 cells closing a void. The yellow line is the boundary of the 3D microvoid substrate. c, 3D reconstruction of z-projection images and the corresponding y-z and x-z sections of cells (along the white dotted lines) with immunofluorescence staining for F-actin (red) and nuclei (blue) in a 3D microvoid during void closure. F-actin was under a stretched state according to the morphology of the nuclei, which were in an elongated form. d, Decay of the area over time for different void sizes ranging from 100 to 300 μm diameter. The red line corresponds to a void size of 100 μm, blue corresponds to 200 μm, and black corresponds to 300 μm. e, SEM of cells partially closing a void from the 30° tilted top view. The scale bars are 100 μm (b) and 20 μm (c).

### Artificial gap closure of stromal cells mediated by purse-string mechanism

During the gap closure, stromal cells were well aligned morphologically at the periphery of advancing cells. We allowed the cells to partially close the gap and immunostained with anti-phosphorylation myosin light chain 2 (PMLC) antibody to visualize actin filaments in stress fibre forms. To our surprise, stress fibres accumulated and were well aligned along the gap edges (Fig. 2**a** and **b**). During gap closure, the boundary profile formed by the stromal cells showed a smooth landscape (Fig. 2**c-e**) rather than a tortuous morphology typical of lamellipodia. Hence, we hypothesized that fibroblasts could close void in free spaces by utilizing actin stress fibres, similar to the epithelial purse-string mechanism, which has never been reported for gap closure by stromal cells. We compared the 2D wound healing process between stromal and epithelial cells by using the same substrate. Unlike epithelial purse-string contraction and its subsequent shrinking of the wound periphery (Supplementary Fig. 5**a** and **d**), 2D mesenchymal gap closure exhibited a tortuous boundary formed by heterogeneous and dynamic lamellipodia of migrating leader cells (Supplementary Fig. 5**b** and **e**). While epithelial cells exhibited highly coordinated movement towards the wound centre, mesenchymal cells closed the wound by only a few highly motile leader cells that migrated towards the wound centre (Supplementary Fig. 5**c**). The directional persistence of the two cell types showed significant differences where the stromal cell gap closure on the 2D substrate was dominated by randomly activated cell crawling rather than uniform purse-string contraction. The degree of tortuosity, defined as the ratio of the contour peripheral length to the geometrical length per unit arc (unit angle = 30°), increased persistently until the gap was nearly closed by mesenchymal cells while epithelial wound showed a low degree of tortuosity maintained over time (Supplementary Fig. 5**f**). These differences in wound morphology change likely stem from the distinctive migratory patterns of the two different cell types near the wound edge. According to the results of PMLC staining, the expression was random everywhere, suggesting no purse-string contraction on the 2D planar substrate (Supplementary Fig. 5**g**). This result is also consistent with a previous report on the 3D wound healing process^16^. When stromal cells are treated with pharmacological agents such as blebbistatin or Y27632 that interfere with actomyosin contractility and its upstream Rho-ROCK pathway, the wound closure rate in the 2D wound healing process was not significantly affected. This finding confirms that the wound closure by stromal cells was not primarily performed by actomyosin-driven purse-string-like closure (Supplementary Fig. 6). Therefore, contrary to the gap closure of epithelial cells on 2D planar substrates, which typically depends on both actively migrating cells with lamellipodia and the purse-string contraction by supracellular actin cables assisted by proliferating cells^21^, the fibroblasts were not expected to employ purse-string contraction in the 3D microvoid closure. However, unlike the tortuous boundary observed in 2D mesenchymal gap closure, the mesenchymal cells displayed a smooth boundary in the 3D microvoids. Taken together, these results indicate that fibroblasts align themselves along the periphery to close the gap by purse-string-like contraction and utilize the cortically aligned actin cables along the edge, which was only previously observed for epithelial cells on 2D planar substrates^7,9,12,22^. It should also be noted that we did not observe gap closure and bridge formation by epithelial cells (HaCaT and Kera-308) in 3D microvoids.

**Fig. 2.**
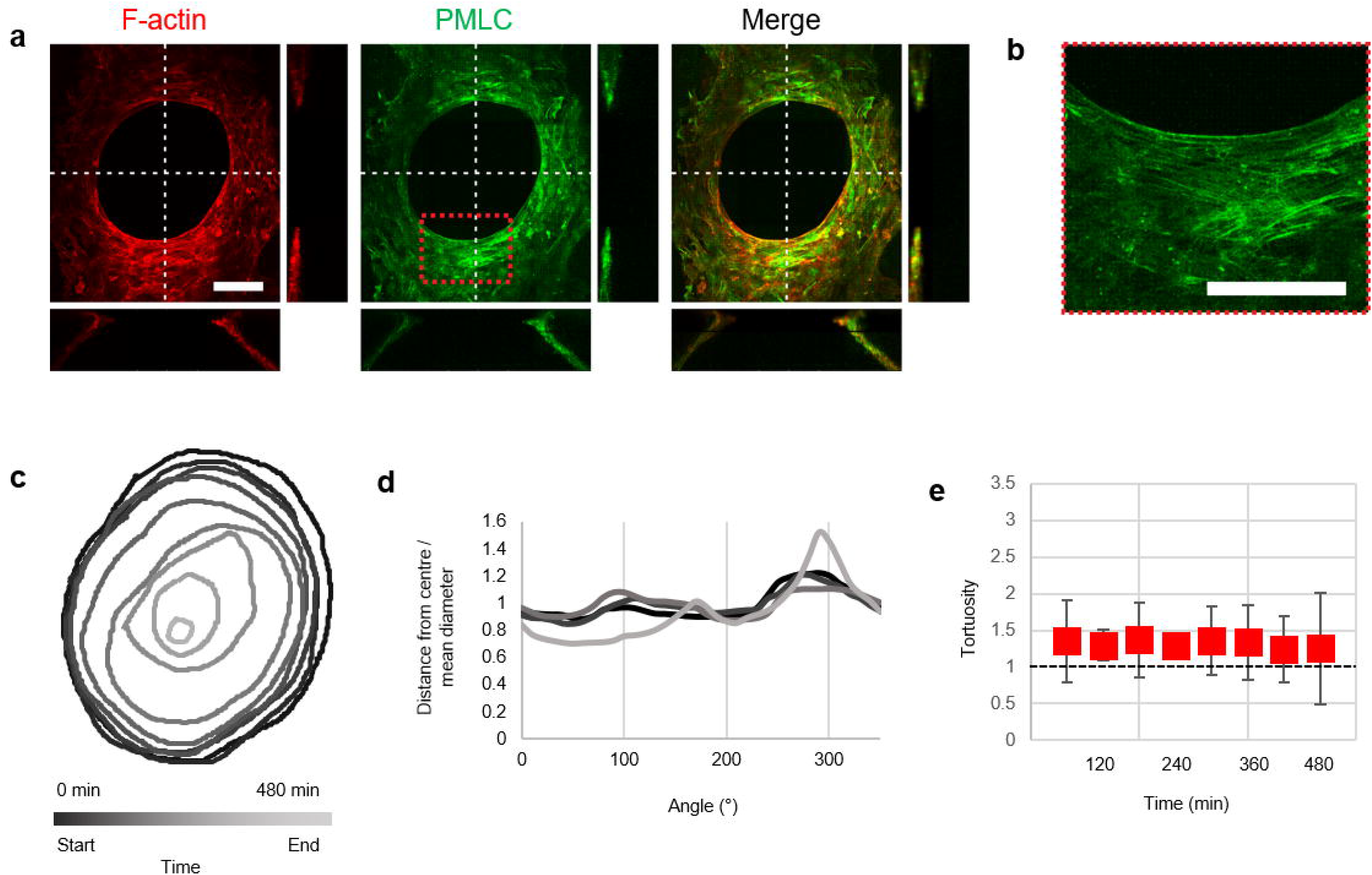
Fibroblast cells close voids via purse-string contraction. **a,** Stromal cells can close a void by employing purse-string contraction, which has been mostly observed only in epithelial cells. 3D reconstruction of the z-projection of confocal images and the corresponding y-z and x-z sections of cells (along the white dotted lines) stained for F-actin (red) and phosphomyosin light chain (PMLC, green) during void closure. **b,** The highlighted box from (a) shows well-aligned PMLC at the periphery of the void as evidence of the purse-string mechanism. c, Shape changes of the void boundary during the course of void closure. **d-e,** The tortuosity of stromal cells migrating in the 3D microvoid observed over 480 min. The smooth boundary indicates good alignment of actomyosin contractility at the periphery of the void and the existence of purse-string contraction during void closure of 3D microvoids. The tortuosity was calculated as Tortuosity = L/L0 (L=total length of the curve of the unit angle (here, 60° is selected as a unit angle), L0 = length of the curve when the whole wound boundary is in a circle with a mean diameter). The scale bar is 70 μm.

### Gap closure of stromal cells in 3D microvoids relies on myosin II-mediated purse-string contractility

To further confirm the purse-string mechanism of fibroblasts during the void closure, we suppressed myosin II contractility by adding 25 μM blebbistatin to the cell culture media. When myosin II activity was suppressed, there was no bridge formation, suggesting the importance of the myosin II-mediated contractility in initiation of gap closure by fibroblasts (Supplementary Movie 4). Moreover, the inhibition of myosin II after a partial gap closure showed a cell retraction to the initial opening size (Fig. 3**a** and Supplementary Movie 5). During the 9 hr observation, the cells retracted, enlarging the gap area to nearly 567% (Fig. 3**a** and **c**) due to the loss of pre-tension in the monolayer by blebbistatin altering the structural integrity of the stress fibres (Fig. 3**c** and **d**). However, there was little alteration of nucleus elongation in the system (Fig. 3**d**), unlike the case when a 2D flat surface was treated with blebbistatin, for which the nuclei recovered to a rounded morphology^23^.

**Fig. 3.**
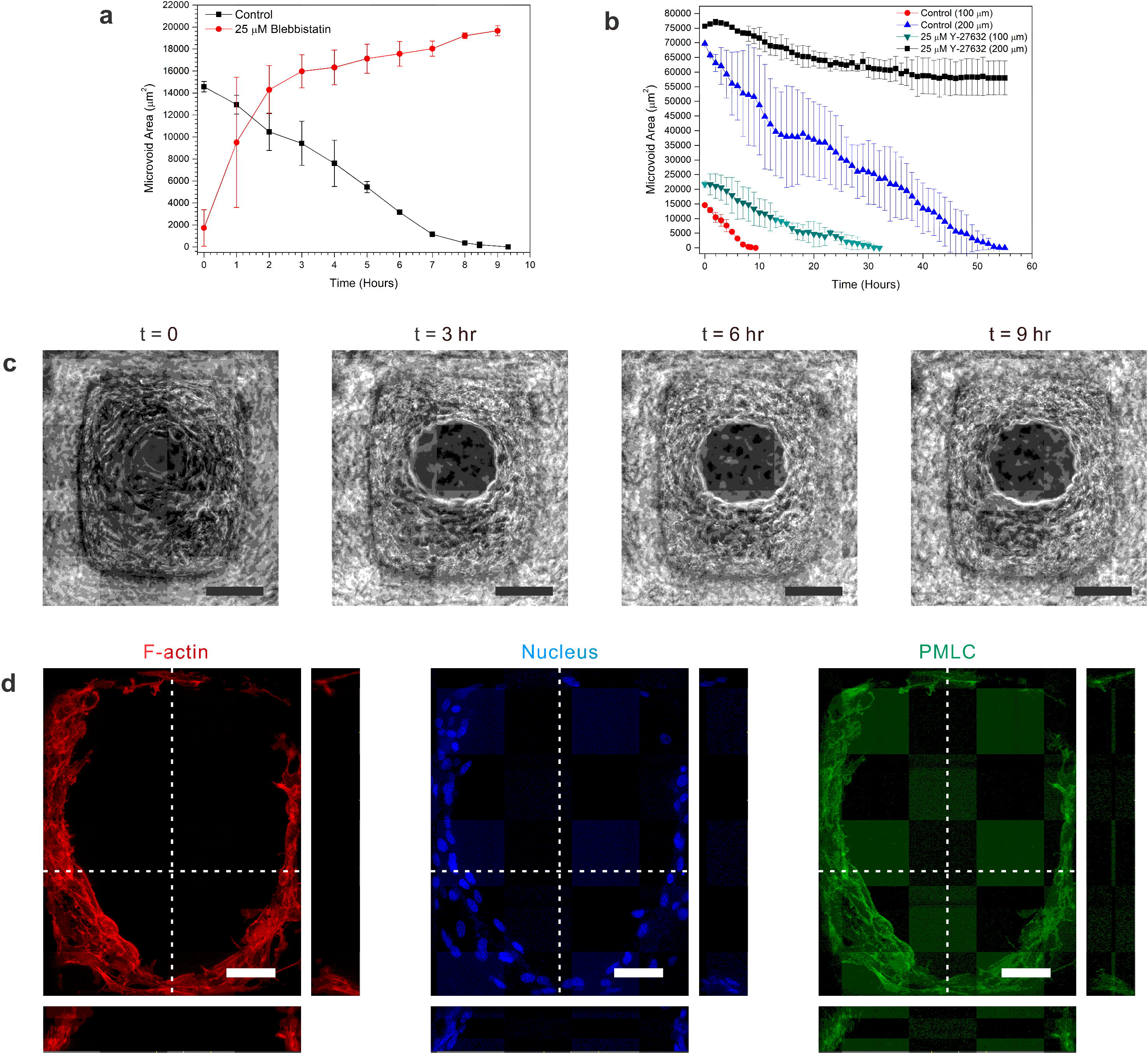
Void closure is mediated by the contractility of myosin II and Rho-associated protein kinases (ROCKs). **a,** Cells were retracted when treated with 25 μM blebbistatin. The retraction of cells reached ~520% within 5 hr and was saturated at this time point. **b,** The independence of Rho-associated protein kinases (ROCKs) for smaller voids, although ROCKs were required to efficiently close large voids. **c,** Phase contrast microscopy images of the retraction of cells treated with 25 μM blebbistatin. **d,** Immunofluorescence staining of F-actin (red), nuclei (blue) and adherens junctions (green) after myosin II contractility was inhibited. Blebbistatin was found to alter cells’ F-actin morphology from a stretched state to a relaxed state. The scale bars are 50 μm (**d**) and 100 μm (**c**).

We also investigated the role of stress fibre-mediated intracellular tension during gap closure by inhibiting Rho-associated protein kinase (ROCK) with 25 μM Y-27632 before the formation of cell bridges. The results showed that cells were still able to close a smaller gap (100 μm diameter), although the closing rate was much slower than that of control (Fig. 3**b** and Supplementary Movie 6). Additionally, retarded closure of the gap was more pronounced in large gaps (>200 μm) (Fig. 3**b**, Supplementary Fig. 7**a** and Supplementary Movie 7). The effect of ROCK inhibition was observed at the initial gap closure, evidenced by the slow slope of the closure (Fig. 3**b**). We also noted that, unlike the case of myosin II inhibition, ROCK inhibition relaxed neither the stress fibres completely around the edge of the gap nor the elongated nuclear morphology (Supplementary Fig. 7**b**-**e**). Taken together, these data show that both myosin II and ROCK play a crucial role in the circumferential contractility of actin cables during gap closure. To further assess, we then treated cells with 1 nM calyculin A, a serine/threonine protein phosphatase inhibitor that increases myosin contractility to examine how the myosin contractility affect the void closure. With the myosin II enhancer, calyculin A, the rate of closure was greatly increased, as shown in Supplementary Fig. 8, with a mean velocity of 11.42 ± 0.94 μm/hr, which was almost two-fold that of normal gap closure. Our results demonstrated the importance of myosin II-mediated contractility of actin bundles during the closure of 3D microvoids.

Furthermore, our results confirmed that 3D microvoids are closed by a purse-string mechanism that relied heavily on the contractility of myosin II.

### Intercellular adherens junctions are crucial during gap closure of microvoids

Sheet migration, another mechanism for the proper gap closure, requires stable cell-cell junctions. When assessed via immunofluorescence of β-catenin, an adherens junction molecule, we confirmed a strong expression of β-catenin, co-localized with actin, during gap closure on 3D microvoids (Fig. 4**a**). To examine the role of adherens junctions, we treated cells with a media containing low Ca^2+^ (~0.03 mM calcium chloride) to weaken the strength of intercellular adherens junctions. First, we verified the inhibition of adherens junctions by immunofluorescence staining of cells cultured in low Ca^2+^ and confirmed that adherens junctions (β-catenin) were downregulated (Supplementary Fig. 9). Next, the same set of experiments on the microvoids showed that the cells were unable to establish cell bridges under low Ca^2+^ conditions (Supplementary Movie 8). In another test, when we allowed the cells to partially close the gap and then changed the media to low Ca^2+^, the cells neither continued the gap closure nor retracted back (Fig. 4**b** and Supplementary Movie 9). The lack of retraction implied that low Ca^2+^ did not compromise intracellular actin stress fibres despite the fact that the cells failed to follow the cells at the periphery to advance forward for proper closure. These data suggest that intercellular adherens junctions are a critical part of gap closure on microvoids.

**Fig. 4.**
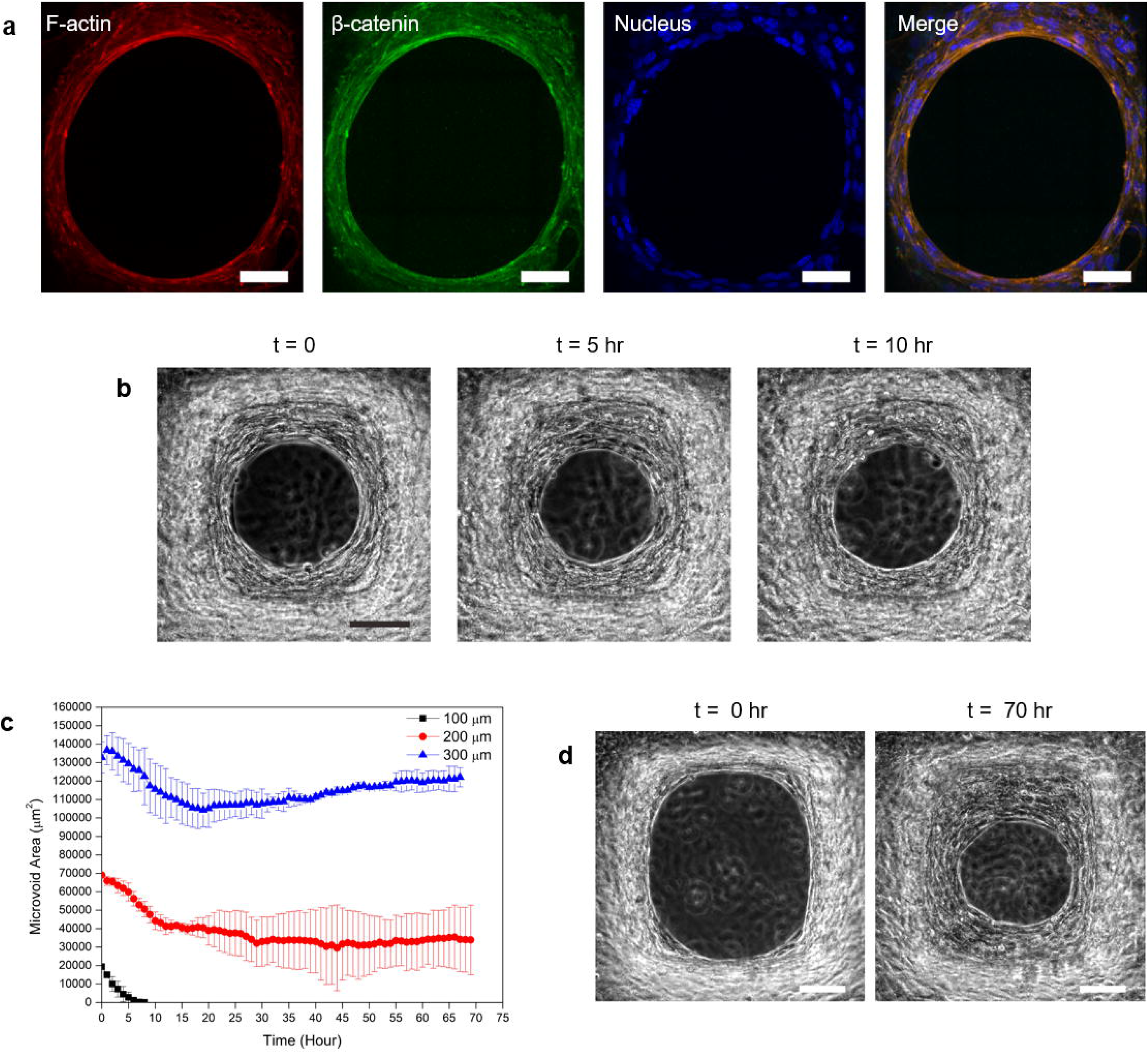
The roles of intercellular adherens junction and cell proliferation in 3D microvoids. **a,** Immunofluorescence staining of F-actin (red), nuclei (blue) and β-catenin (green) during void closure. The presence of β-catenin suggested that fibroblast cells close the void through robust adherens junction. **b,** Inhibition of intercellular adherens junctions using low Ca^2+^ discontinues void closure. **c,** Plot of the role of cell proliferation in void closure with different void sizes. The efficiency of void closure with cell proliferation inhibition was equal to that of normal void closure for smaller voids, although larger voids were not fully closed, indicating that cell proliferation is essential for efficient void closure. **d,** Phase contrast live-cell microscopy images of cells treated with 10 μM mitomycin C before formation of the bridge (*left*) and after 70 hours of void closure (*right*). The scale bars are 25 μm (**a**) and 100 μm (**b, d**).

### Efficient gap closure requires cell proliferation for larger gaps

For further assessment, we investigated the role of cell proliferation as the driving force of gap closure. When cell proliferation was inhibited by treatment with 10 μM mitomycin C, a smaller gap (100 μm) was closed completely within 10 hours before the proliferation effect became important, given that the typical doubling time of fibroblasts would be at least a day (Fig. 4**c**). On the other hand, the larger gaps (200 and 300 μm), which would take longer than the doubling time, failed to fully close (Fig. 4**c** and **d**). For 300 μm gaps, the void region became even larger with time, indicating that the intercellular pressure in the monolayer due to crowding was lost by the suppression of proliferation. We suggest that the initial gap closure was accomplished by the purse-string mechanism of the existing cells along the periphery, which would then be assisted by the proliferating cells in the bulk region. Cell proliferation contributed significantly to efficiently closing larger gaps, not significantly affecting the initiation of the gap contraction.

### Zip-up gap closure mediated by directional purse-string contraction for 3D U-shaped microgaps

Wounds are not always circular in shape. We fabricated an open contour of a microgap with a U-shaped edge, as described in Supplementary Fig. 10. As fibroblasts are seeded on the flat base of a 3D U-shaped microgap substrate, they form a bridge-like, free-standing cell layer covering the gap, analogous to a “zip up” process. As in the microvoids, the cells established a bridge that was initiated at the κ_corner_ in the rounded edge and subsequently closed the gap in a free-standing manner without any supporting matrix (Fig. 5**a, b** and Supplementary Movie 10). During the initiation of bridge formation, we observed some retraction of the individual cells, indicating that adherens junctions of the collective cells were crucial for robust cell bridge formation (Supplementary Fig. 11**a**-**h**). We monitored stable cell bridge formation at 5 hr post-seeding by time-lapse microscopy (Fig. 5**c**). During the course of cell migration, fibroblasts created robust bridge formation along the 3D U-shaped microgap, as confirmed by SEM (Fig. 5**d**). Cells on the microgap formed thick actin cables that spanned over multiple cells along the edge at t = 85 hr, as shown in Fig. 5**e**. These cells did not exhibit any lamellipodia at the advancing front, but instead, they were pulled closer by contraction of the actin cables formed along the periphery of the cells (Supplementary Movie 11). At t = 85 hr, the gap was filled by a cell sheet composed of vertically stacked multiple cell layers, as shown in the cross-sectional image through y-z. The cross-sectional image through x-z shows that the cells travelled over the edge to the underside, forming a continuous support across the gap.

**Fig. 5.**
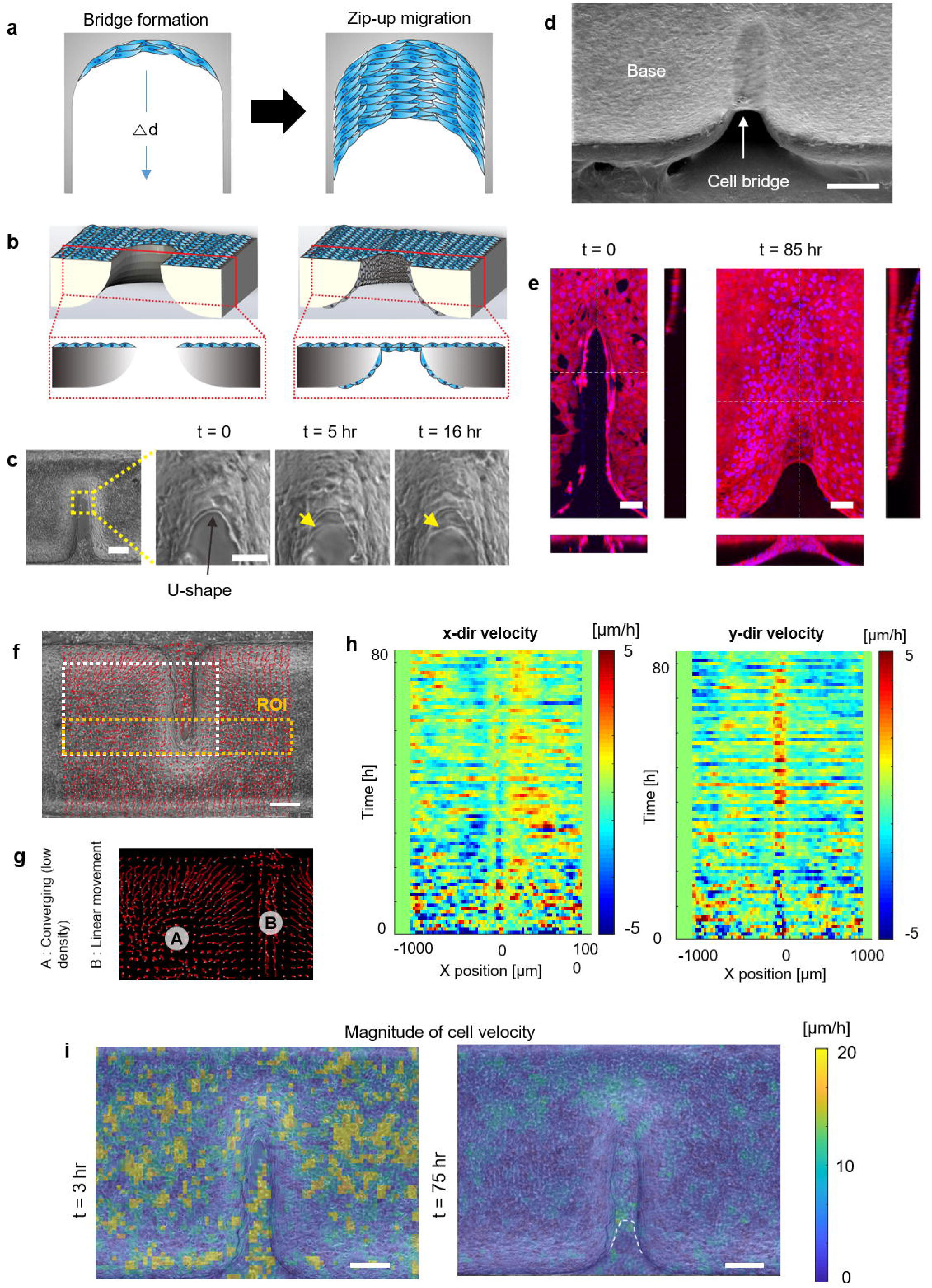
Zip mechanisms of gap closure of 3D linear microgaps. **a** and **b,** Diagram of the top view (**a**) and tilted view with a cross-section (**b**) of initial cell bridge formation and zip gap closure of a 3D linear microgap. Δd is the displacement. **c,** The initiation of cell bridge formation observed through a live-cell imaging confocal microscopy. Five hours after cell seeding, cells formed stable adherens junctions (arrows) initiated at the U-shape structure. Before establishing robust adherens junctions, retractions were also observed (Supplementary Fig. 11). **d,** SEM representation of cell bridge formation three days after cell seeding. **e,** 3D reconstruction of the z-projection of confocal images and the corresponding y-z and x-z sections of cells (along the white dotted lines) stained for F-actin (red) and nuclei (blue) before bridge formation at t = 0 (*left*) and after zip-up gap closure at t = 85 hr (*right*). **f-i,** Kinematic changes in cell migration during the zip-up mechanism. **f,** Trajectories of cell migration during zip-up gap closure. **g,** The highlighted (white) boundary in (**f**). Cells on the microgap region clearly migrate with linear movement along the microgap analogous to zip-up migration. Longer lines indicate faster motion of cells, and straight lines represent more persistent motion. **h,** The kymograph of x-directional velocity in the bridge formation condition. Each row corresponds to a region of interest (ROI). **i,** Map of the speed of cellular movement according to the time during wound healing on the U-shape structure. The white dotted line indicates the cell bridge. The scale bars are 100 μm (c *left* and e), 50 μm (c *right*), 250 μm (d) and 200 μm (f and i).

Similar to the gap closure on the microvoid, we found that the zip-up gap closure was also mediated by purse-string contraction, as evidenced by PMLC expression. The results revealed well aligned PMLC at the periphery of the advancing cells (Supplementary Fig. 12).

Additionally, blebbistatin treatment after the adhesion stage showed no bridge formation. To investigate whether the collective motions of the cells on the flat base contributed to the zip-up gap closure, we plotted the trajectories of cell motion over 80 hours. We noticed faster and persistent movement of cells near the far edges of the gap, indicating the continuous movement of cells over the gap edges, however, cells along the gap edge showed almost no motion or converging motion to the gap, suggesting that the zip-up gap closure is not influenced by cells on the flat base along the periphery of the gap edges (Fig. 5**f** and **g**). Once the bridge is formed, the cells move persistently along the long direction of the gap. This persistent zipping motion can be clearly observed, as shown by the positive red vector in the y-directional velocity at position 0, the centre of the gap beyond 30 hours of time (Fig. 5**h**). In the x-directional velocity kymograph, no feeding of cells was observed towards the gap within the ROI. In particular, once the bridge started to form and grow between gaps for approximately 30 hours, a laterally outward motion was observed with respect to the gap axis. The magnitude plot of cell velocity at t = 3 hr shows sporadic hot spots that are not located near the edge (Fig. 5**h**). The finite movements detected inside the gap before the gap was filled at early time points are due to the fibroblasts on the bottom of the chamber, which we have shaded out in the image to avoid confusion. After the gap was closed (t = 75 hr), overall migration was suppressed except for those that moved on the bridge at the centre of the gap (Fig. 5**i**). These results suggest that zip-up gap closure mostly involves cells that migrate on the microgap and not from the base. In conclusion, we propose that the purse-string contraction is a driving force for closing U-shaped 3D microgaps via a zip-up mechanism.

### Geometrical size effect on purse-string gap closure

To further understand how stromal cells close gaps of different shapes, we investigated the dependence of efficient gap closure on the curvatures of the advancing cells (κ_advancing cells_). Given that substrates have the same κ_corner_ (0.02 μm^−1^), the bridges were formed irrespective of the overall gap geometry (Fig. 6**a**). After the initial formation of the bridge at the corner with curvature κ_corner_, the advancing cells established a finite curvature by the aligned cells, κ_advancing cells_, both on the open contour 3D U-shaped microgap and in the 3D microvoid (Fig. 6**b** and **c**).

**Fig. 6.**
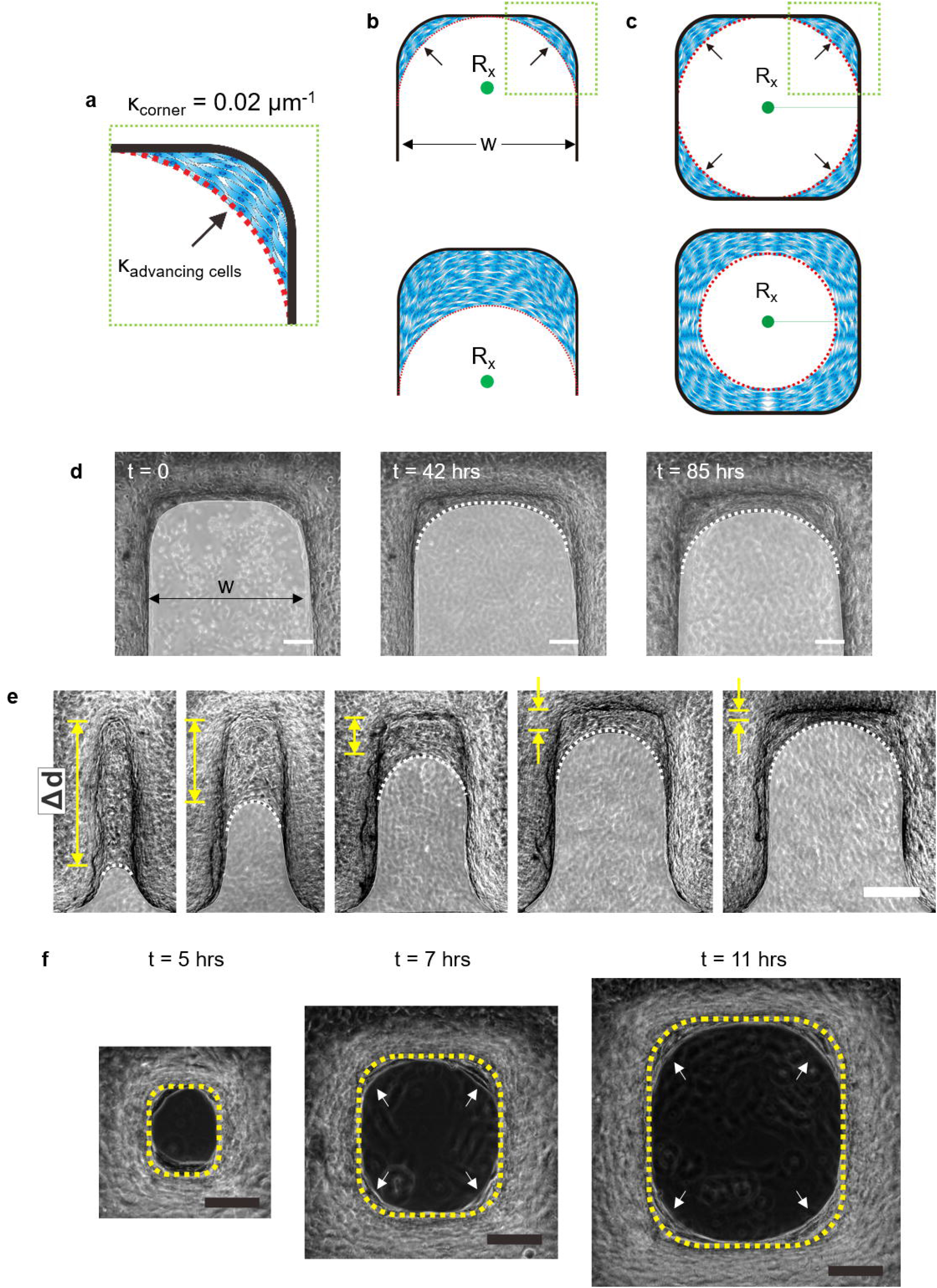
The efficiency of gap closure relies on the curvature of advancing cells. **a,** Illustration of the formation of the κ_advancing cells_ indicated by the arrows after initiation of bridge formation at the κ_corner_. **b** and **c,** Over time, the cells formed a curve shape (**b**) or a circular shape (**c**, red dot lines) in 3D linear microgaps and microvoids, respectively. R_x_ is the radius of the curvature generated by the cells. **d,** Time-lapse microscope images of cells that formed curves in 3D linear microgaps with larger widths (w: 500 μm). White dotted lines are the κ_advancing cells_. **e,**Δd for different gap widths at 85 hr. White dotted lines indicate κ_advancing cells_ during zip-up gap closure. A larger κ_advancing cells_ corresponds to a smaller Δd. **f,** Time-lapse images of κ_advancing cells_ (arrows) formed after initial bridge formation with various 3D microvoid sizes (100-300 μm). To effectively close the void, the size of the κ_advancing cells_ assembled by the cells should be higher than 0.0066 μm^−1^. Though the gaps and voids have various sizes, the curvature of the PDMS corners (κ_corner_ = 0.02 μm^−1^) is equal. Thus, formation of the bridge can be initiated for all sizes of voids. The yellow dotted lines are the boundaries of the PDMS substrate. The scale bar is 100 μm.

As long as κ_corner_ was kept the same, the gaps of any size could initiate bridge formation whose curvature was κ_advancing cells_ at κ_corner_, although more time was required for a larger gap width (w) to form κ_advancing cells_ (Fig. 6**d** and **f**). However, during zip-up gap closure, the gap width determined the displacement (Δd) and the κ_advancing cells_ (Fig. 6**e** and Supplementary Fig. 13). The quantitative analysis provided in Supplementary Fig. 14 showed that after 85 hr, the Δd during bridge formation was higher in the 100 μm gap (273 ± 78.6 μm) with a velocity of 3.41 ± 0.98 μm/h. The lowest Δd was observed in the 500 μm gap with significantly less Δd (32.16 ± 25.73 μm) and much lower velocity (0.40 ± 0.32 μm/h). These results indicate that the larger the width of the gaps is, the smaller the κ_advancing cells,_ which therefore results in a smaller Δd. Our results also revealed that the velocity was proportional to Δd (Supplementary Fig. 14). These findings confirmed that gap closure succeeded when κ_advancing cells_ was higher than 0.0066 μm^−1^.

Consequently, when we tested a larger void with a diameter of 400 μm (κ_advancing cells_ = 0.005 μm^−1^), cells indeed formed bridges at the κ_corner_, but the closure was incomplete and inefficient. These results suggest that the κ_advancing cells_ generated by the cells after the initial bridge formation was the determinant of the void closure by the stromal cells via purse-string contraction.

We next questioned whether cells would still form bridges to eventually close the gaps if κ_corner_ was removed. To test this, we maintained the gap widths but replaced the U-shaped end structure with a circular end with a radius larger than the gap width. We found that there was no formation of bridges, indicating no gap closure except for a 10 μm gap width (Supplementary Fig. 15). These results imply that the structural geometry of the substrate is crucial for recruiting cells and consequently forming robust intercellular adherens junctions to initiate bridge formation. Furthermore, these results demonstrate that stromal cells initiate purse-string contraction when a critical local curvature, κ_corner_, is present to act as a biophysical cue, leading to collective migration to facilitate zip-up closure of 3D U-shaped microgaps. It should also be noted that in epithelial gap closure on a non-adherent 2D planar substrate, the critical diameter to close the gap was reported to be ~150 μm due to the sole dependence of epithelial cells on purse-string contraction with no involvement of active cell crawling^9^. Interestingly, however, our study shows that stromal cells may cover a 3D U-shaped microgap with a larger gap width that is up to two-fold greater than the critical gap closure size of epithelial cells on non-adherent substrates. This feature may be due to the higher focal adhesion of stromal cells over epithelial cells at the onset of bridge formation^24^.

## Discussion & Conclusion

Most research on collective cell migration has addressed how cells close the gap on the 2D planar substrate, which does not mimic the *in vivo* microenvironment of more complex geometrical cues^25–27^. Our results revealed that stromal cells close 3D deep microvoids, and they featured a remarkably similar closing mechanism to that of epithelial cells on 2D planar substrates. Purse-string contraction, which had been previously believed to be a *bona fide* epithelial wound closure mechanism, was employed by NIH 3T3 mouse fibroblasts to close deep microgaps by forming a free-standing bridge over the void. Stromal cells in 3D microvoids aligned themselves along the periphery, forming circumferential actomyosin cables, which eventually led to the closure of the gap by purse-string contraction via cortically aligned actin cables along the edge (Fig. 2**a**). During the gap closure, no lamellipodial extensions were formed at the leading edge of the cells, which contradicted previous reports^9^. Purse-string mechanisms of stromal cells on 3D microvoids were further proven by confirming the existence myosin II contractility including ROCK and the reliance on intercellular adherens junctions. Unlike previous reports, gap closure by stromal cells in our system was shown to heavily depend on the contractility of myosin II, as evidenced by the failure to form a bridge when myosin II contractility was inhibited (Supplementary Movie 4). The retraction of the cell sheet was observed if the contractility of myosin II was inhibited while the closure progressed (Fig. 3**a** and **c**). We reasoned that this retraction was due to a loss of tension in the actin cables in the advancing cells of the periphery, causing shrinkage of the cells. The loss of tensional stress propagated very quickly to cells in the inner layer to cause massive shrinkage (Fig. 3**d**).

Additionally, when ROCK was inhibited, closure took place; however, the closure rate was significantly decreased by two-fold compared with normal gap closure (Fig. 3**b**). Taken together, our results showed that intracellular contractility of myosin II, including ROCK, was required, directly suggesting the involvement of purse-string contraction during void closure.

Stromal cells are known not to maintain stable cell-cell junctions. Instead, they mostly involve transient cell-cell adhesions^28–30^. When stromal cells are seeded on a non-adherent substrate, cells cannot succeed in bridge formation, presumably due to the lack of stable adherens junction proteins^14^. In our study, we found that intercellular adherens junctions played an important role in closing the gap. As shown in Fig. 4**a**, β-catenin is indeed present during the course of gap closure. The reliance of gap closure on the intercellular adherens junction was verified when we inhibited adherens junction protein before the gap closure stages. We found that bridge formation was not initiated when intercellular adherens was inhibited (Supplementary Movie 8). Additionally, when we allowed the cells to partially close the gap and inhibited adherens junction protein, to our surprise, the cells neither retracted back nor advanced forward to close the gap (Fig. 4**b** and Supplementary Movie 9). These results proved that the intercellular adherens junction of stromal cells was critical and essential in closing 3D microvoids. These results also imply that the closure of stromal cells on 3D microvoids was driven by purse-string contraction. We observed that the assembly of actomyosin cables that contributed to the purse-string contraction at the wound edges involved adherens junctions, similar to the mechanism of epithelial cells^31–33^. Therefore, we suggest reconsidering the role of adherens junctions in regulating stromal cells as a crucial parameter for determining gap closure, as observed in this report.

We demonstrated that by changing microvoids into open contour 3D microgaps, gap closure was achieved by a zip-up mechanism and was still mediated by purse-string contraction (Supplementary Fig. 12). Zip-up gap closure indicated that the mediation of purse-string contraction can be established not only in the circular-ring, as in the microvoid case, but can also be mediated in half-circular-like gaps in open contour microgaps. Therefore, we suggest that stromal cells may establish purse-string contraction at the onset of bridge formation initiated at the κ_corner_ of the substrates. Our results confirmed that stromal cells exhibit epithelial-like behaviours, such as purse-string contraction and dependence on robust intercellular adherens junctions, which contradicts previous results. The closure mechanisms of stromal cells described in this study essentially rely on the contractility of myosin II, including ROCK. Additionally, well-aligned multicellular actomyosin structures at the periphery of the microvoid or microgap exert a contractile force for closing the gap. These findings help to further elucidate the role of stromal cells *in vivo* and their contribution to wound healing and tissue regeneration.

## Methods

The 3D microvoid substrates were prepared using a custom-engineered stainless steel mould. The mould of the substrate was designed by using AutoCAD software, and the pattern was transferred for mask fabrication (Supplementary Fig. 1**a**). Once the patterned mask was printed out and placed onto stainless steel, the exposed surfaces were etched out using chemical etching. The fabricated stainless steel mould was then coated with polytetrafluoroethylene (PTFE) using deep reaction-ion etching with a thickness of 10 nm to degrade the adhesion between PDMS and the stainless steel mould during the peel-off process. 3D microvoids were created from a curing agent and the base of a PDMS kit (Sylgard 184, Dow Chemical) by mixing the two components thoroughly at a ratio of 1:10 (curing agent:base). The mixed PDMS solution was then poured into a stainless steel mould and heat-treated for fast curing at 150°C for 10 min by following the technical specifications of *Dow Corning Inc*. After the cured microvoids were peeled off from the mould, they were denoted as 3D microvoids and U-shaped microvoids.

### PDA coating

3-Hydroxytyramine hydrochloride (dopamine hydrochloride, H8502; Sigma-Aldrich) was dissolved in 10 mM Tris buffer (pH 8.5) to a final concentration of 2 mg/mL. PDMS-based 3D microvoid samples were treated by dip-coating in polydopamine (PDA) solution to improve the wettability of the PDMS surface for better cell adhesion. Briefly, PDMS microvoid samples were rinsed with deionized water and then immersed into the PDA solution for 24 hr. Later, they were removed and washed two times using deionized water to remove the excess PDA solution from the PDMS microvoid. The samples were dried using nitrogen gas and stored in a desiccator for cell culture and future application. After coating with PDA, we measured the water contact angle (WCA), which was found to be 63.33°, whereas the WCA of bare PDMS is 121.22° (Supplementary Fig. 2**a** and **b**).

### Cell culture

NIH3T3 mouse fibroblasts (ATCC CRL-1658) were cultured in DMEM supplemented with 10% foetal bovine serum (FBS) and antibiotics (100 U mL^−1^ penicillin and 100 μg mL^−1^ streptomycin; Gibco, Grand Island, NY, USA) under normal culture conditions (37°C and 5% CO_2_). Prior to cell seeding (10^6^ cells/mL), the 3D microvoid sample was fixed on a confocal glass bottom dish (100350, SPL Life Sciences).

### Preparation of pharmacological reagents

Blebbistatin (B0560, Sigma-Aldrich) and calyculin A (C5552-10UG, Sigma-Aldrich) were dissolved in dimethyl sulfoxide (DMSO) at specific concentrations before addition to the cell culture media. Y-27632 (Y0503-1MG, Sigma-Aldrich) was also prepared by dissolving in deionized water. Low Ca^2+^ DMEM was made using non-Ca^2+^ DMEM (21068028, Thermo Fisher) supplemented with 1% FBS and antibiotics (100 U mL^−1^ penicillin and 100 μg mL^−1^ streptomycin). Mitomycin C (M4287, Sigma-Aldrich) was dissolved in deionized water at a concentration of 0.5 mg/mL and used as a cell division inhibitor. The stock solution was diluted in media at a ratio of 10 μL:1 mL (mitomycin C: DMEM). The diluted mitomycin C in DMEM was changed after 4 hours with fresh DMEM for the cell division inhibitor experiments.

### Immunofluorescence staining

For all immunofluorescence staining, cells were fixed with 4% paraformaldehyde for 30 min and then rinsed with 1x PBS three times. Rhodamine-phalloidin (R415, Thermo Fisher Scientific) and 4’,6-diamidino-2-phenylindole (DAPI, Thermo Fisher Scientific) were employed for actin cytoskeleton and nucleus staining, respectively, and images were captured after 30 min of incubation using a confocal microscope (Zeiss LSM 700; Carl Zeiss Micro-Imaging GmbH, Germany). All images were taken in z-stack mode.

β-catenin and PMLC immunofluorescence staining were also performed by fixing the sample with 4% paraformaldehyde and then rinsing with 1x PBS. The samples were permeabilized with 0.2% Triton X-100 in PBS for 10 minutes, washed with 1x PBS two times, and then blocked with 3% bovine serum albumin (BSA) in PBS for 1 hr. The samples were then incubated with primary antibodies as follows: anti-β-catenin primary antibody (E247, Abcam) and phosphor-myosin light chain 2 (Thr18/Ser19) antibody (3674, Cell Signaling Technology) in 1% BSA (1/200 dilution) at 4°C overnight. After the samples were thoroughly rinsed with 1x PBS, they were subjected to treatment with secondary antibodies, including Alexa Fluor 488 goat anti-mouse antibody (A11001, Thermo Fisher Scientific) or Alexa Fluor 488 goat anti-rabbit antibody (A11008, Thermo Fisher Scientific) for 1 hr at a dilution ratio of 1/200. The photographs were captured using a laser scanning confocal microscope.

Immunofluorescence images were processed using Imaris 8.0.1. Briefly, the scanned images were processed by section views for 3D reconstruction or sliced view for image extraction. Section views display three images captured at the same time corresponding to the x-y (right top), x-z (right bottom) and y-z (left) sections. For 3D reconstruction views, section views are preferred for displaying the morphology from all orientations.

### Live cell imaging

Zeiss live cell imaging confocal microscopy (Zeiss live cell confocal, Germany) was used for live cell imaging. After the cells adhered well to the 3D microvoid, the samples were transferred to a live cell imaging incubator and maintained under standard culture conditions of 37 °C and 5% CO_2_ in a humidified atmosphere. Movies were generated by capturing images every 10 minutes and one hour depending on the length of the experiment.

### Area calculations

The acquired time-lapse images from live cell imaging confocal during gap closure were quantified using ImageJ software (V1.46r, imagej.nih.gov/ij/).

### SEM images

SEM images were taken after cells were fixed with 4% paraformaldehyde for 30 minutes and then washed with 1x PBS three times. The PBS was removed, and the samples were mounted in an ESEM (Environment Scanning Electron Microscope, XL-30, Philips) chamber for image scanning.

### PIV analysis

To measure the cellular velocity, particle image velocimetry (PIV) analysis was conducted by using the Image Velocimetry Tool for MATLAB (version: 1.43) software. We used double-pass PIV by the fast Fourier transform (FFT) window deformation algorithm with a first window size of 64·64 pixels and a second window size of 32·32 pixels. Velocity vectors were distributed with a 16-pixel interval (20 μm), similar to the body length of a cell. The trajectory of cells in Figs. 5 and 6 was analysed from the velocity fields. The first location of each trajectory was allocated by dividing the whole region with 32·43 windows. The next location was updated by adding the displacement of each window, which was calculated from the PIV data.

### Statistical analysis

Data were analysed using Origin and GraphPad software, and data were plotted using Origin software. For statistical analysis, data were compared using one-way ANOVA (GraphPad Prism 5, La Jolla, CA). Statistical significance is marked as * *P* < 0.05, ** *P* < 0.01, *** *P* < 0.001 or **** *P* < 0.0001.

## Acknowledgements

This work was supported by the National Research Foundation of Korea (NRF) (NRF-2017R1A2B2007673, NRF-2020R1A2C2007972 and NRF-2019R1A2C2003430), and Creative-Pioneering Researchers Program through Seoul National University.

## Author contributions

I.-S.C. conceived the concept, A.D.C. and I.-S.C. designed the research, A.D.C and R.S. performed experiments, A.D.C and H.T.C. performed modelling and computational tools, A.D.C., H.T.C., S.J.O and R.S. analysed experimental data, A.D.C, K.P., J.H.S and I.-S.C. wrote the paper, and K.P., J.H.S, and I.-S.C. supervised the project. All authors discussed the results and commented on the manuscript.

