## Supplementary Information for "Fibroblasts close a void in free space by a purse-string mechanism"

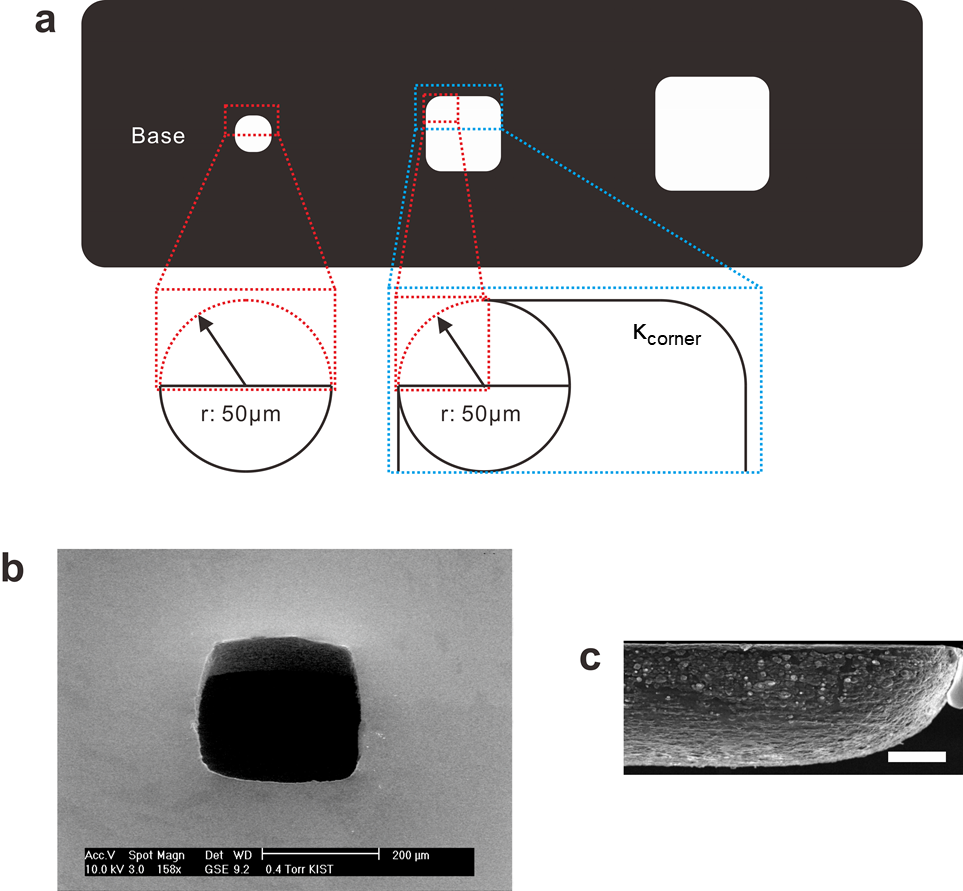

**Supplementary Fig. 1a**. Pattern design of 3D microvoids with varying gap sizes (100-300 µm diameter). Each gap in the microvoids has a quarter circle-like curvature that termed the curvature of the corner (κ_corner_) with a radius of curvature of 50 µm. b, c, Representative scanning electron microscopy (SEM) images of 3D microvoids at 30° tilted and cross-sectional views. The scale bar is 50 µm.

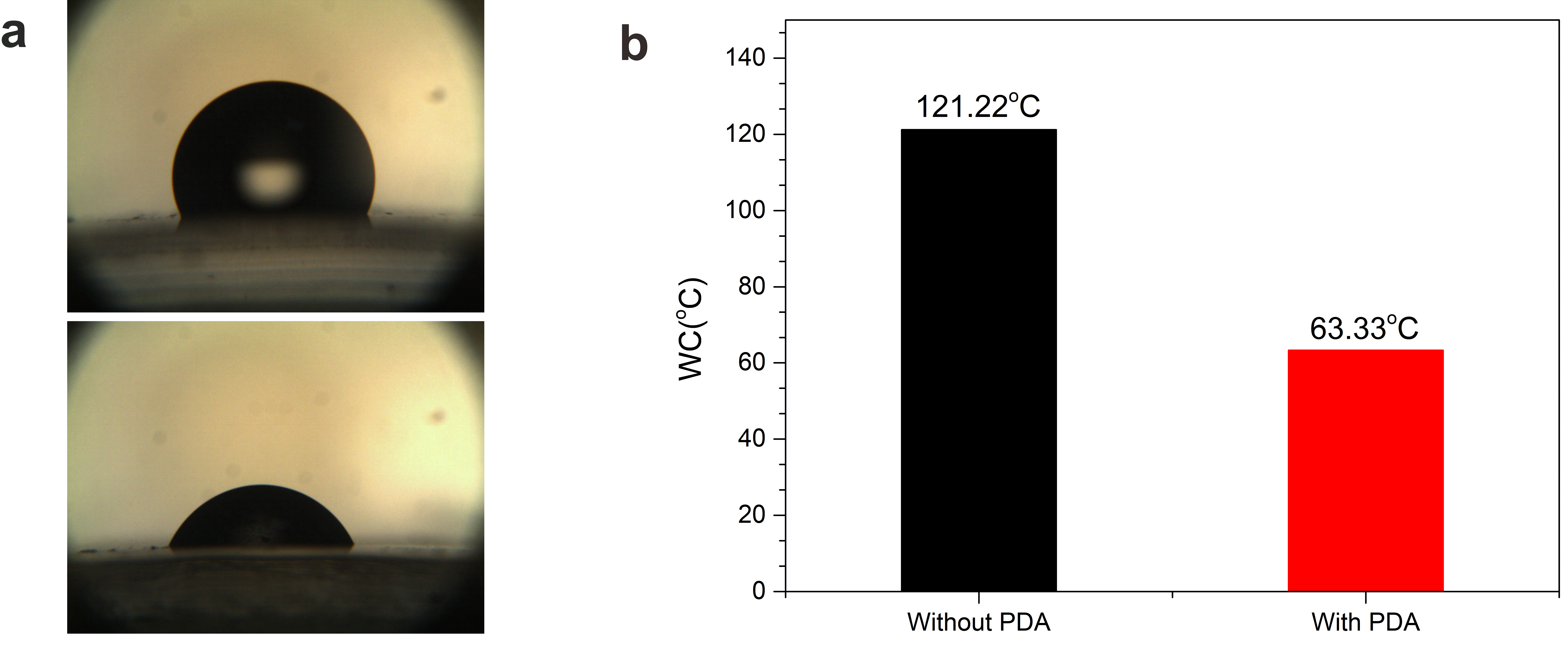

**Supplementary Fig. 2a.** Water contact angle (WCA) before *(top*) and after polydopamine (PDA) coating (*bottom*), b, Quantitative plot of the WCA.

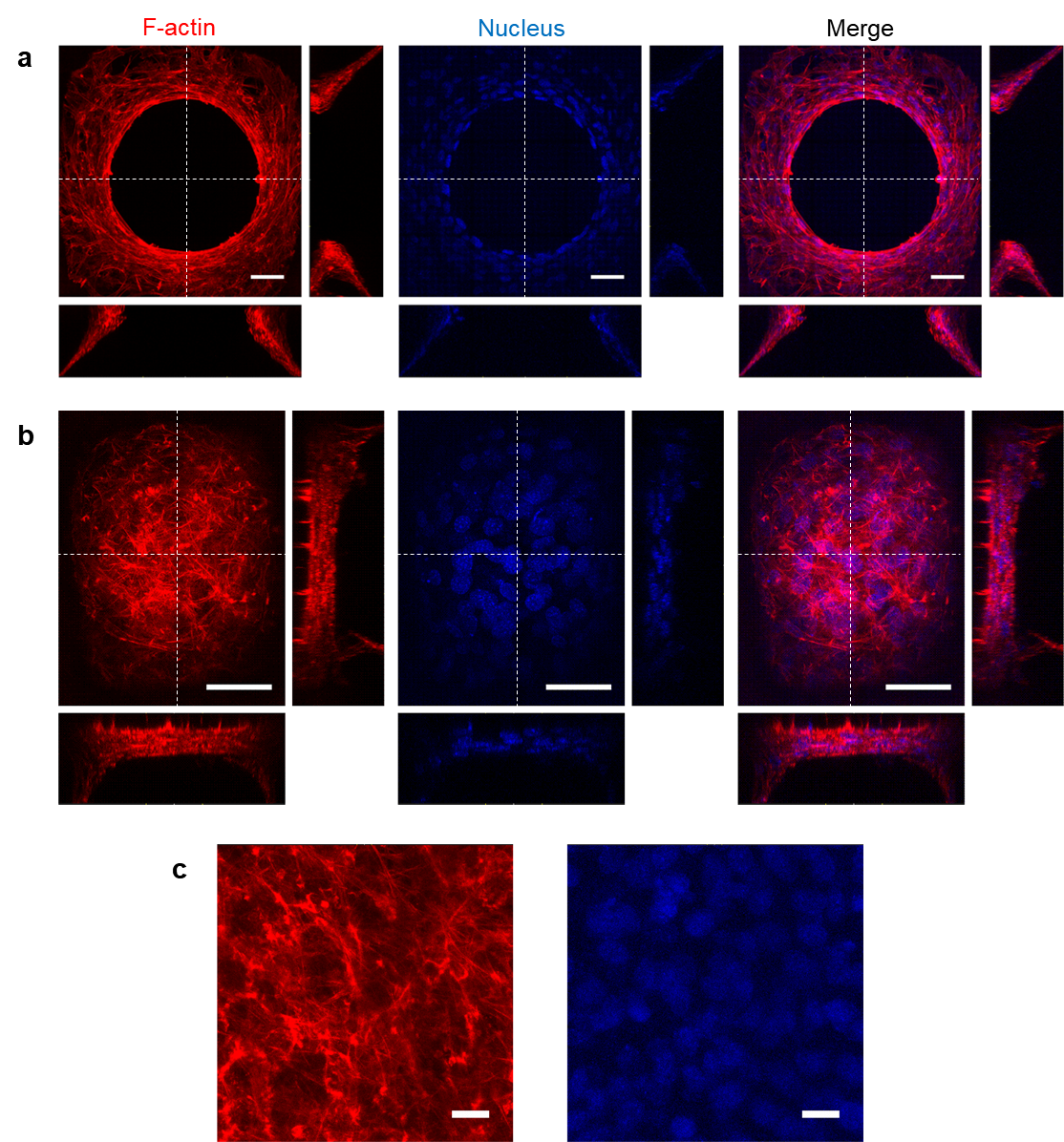

**Supplementary Fig. 3.** 3D reconstruction of the z-projection of confocal images and the corresponding y-z and x-z sections of cells (along white dotted lines) that were immunofluorescently stained for F-actin (red) and nuclei (blue) on rectangular 3D microvoids after void closure. a, Cells formed bridges and partially closed the 3D microvoids. The elongated nuclear morphology indicates that cells on the periphery of the gap are under higher tension. b, With the progression of time, the cells fully closed the void, and the tension decreased, as confirmed by the rounding morphology of the nuclei. c, The cell morphology in the 3D microvoids indicates lower tension. Scale bars are 20 µm (a) and 50 µm (b, c).

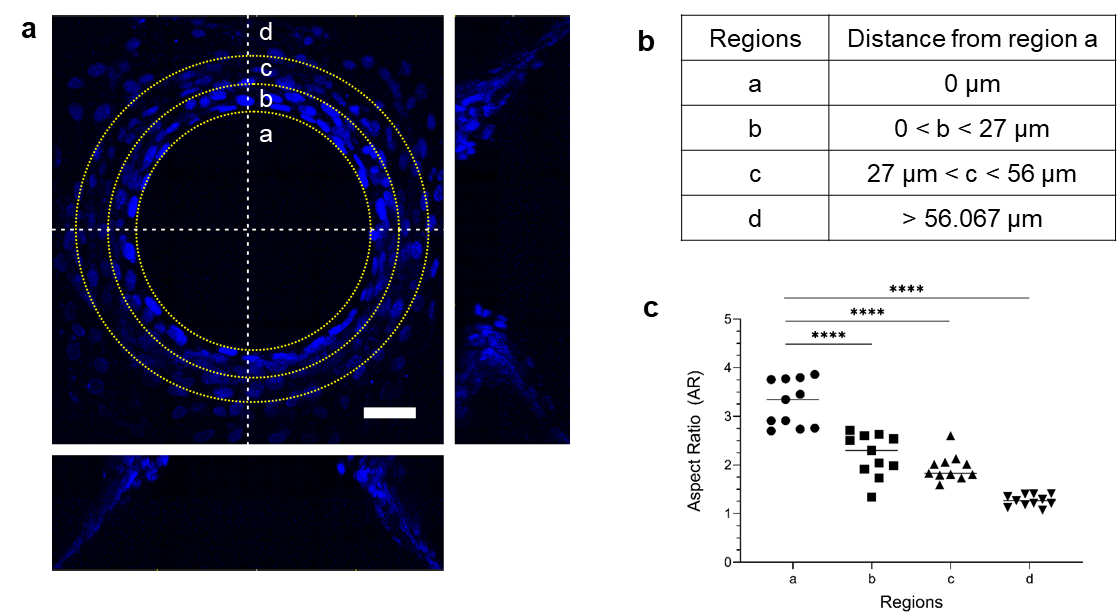

**Supplementary Fig. 4a.** Regional division of the nucleus during void closure on 3D microvoids. The scale bar is 20 µm. b, Table of the distance from region *a*. c, Aspect ratio (AR) of the cells on different regions on 3D microvoids. The cells at the periphery of the advancing migration were under significantly higher tension (region *a*). As the distance into the periphery of the advancing cells increases, the tension distribution decreases until a normal cell tension was reached at distance d. Data were acquired from 2 independent experiments. **** *P* < 0.0001.

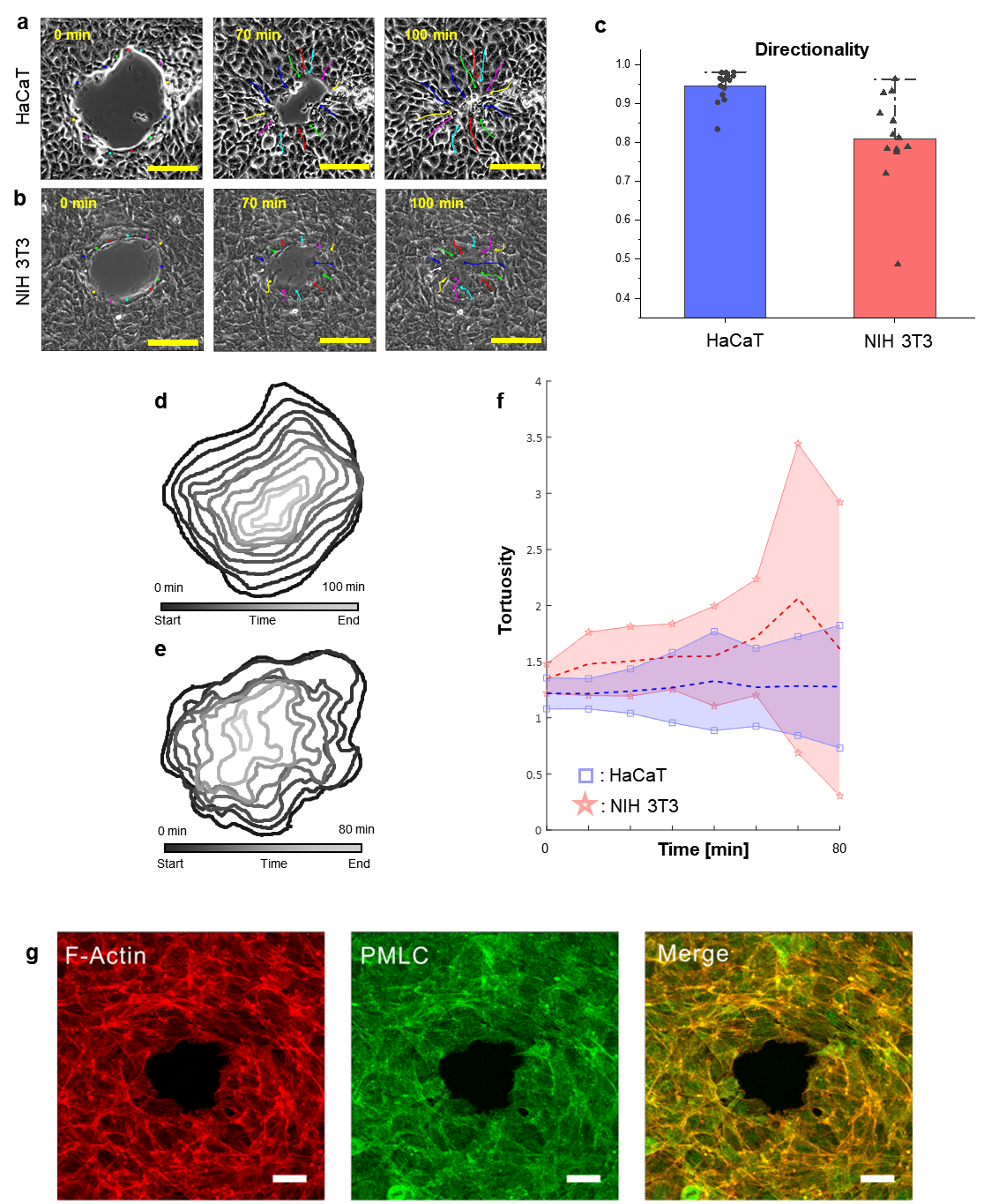

**Supplementary Fig. 5.** 2D planar mesenchymal gap closure involves active lamellipodia rather than purse-string contraction. a,b, Migration trajectories of cells at the wound edge. c, Differential directionality of cell migration between epithelial cells and mesenchymal cells. d, e, Initial shape and trajectories of the wound boundary during wound closure of the HaCaT (d) and NIH 3T3 (e) cell monolayers on a 2D substrate. f, Differential tortuosity of the wound boundary of epithelial cells and mesenchymal cells. g, Immunofluorescence staining of NIH 3T3 mouse fibroblasts on a 2D substrate for F-actin (red) and phosphorylated myosin light chain (PMLC, green) during gap closure. Scale bars are 100 µm (a and b) and 50 µm (g).

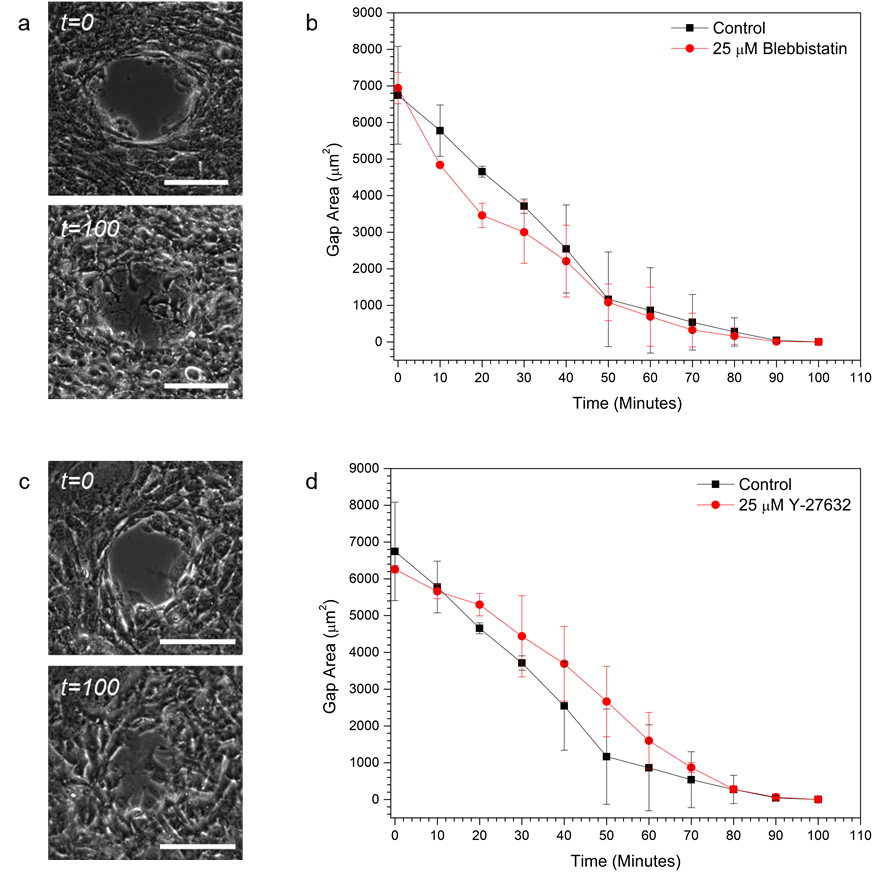

Supplementary Fig. 6. Mesenchymal 2D gap closure is not affected by the inhibition of actomyosin contractility. a,b, Time lapse images of the initial and final closure of the gap when treated with 25 µM blebbistatin. The gap was fully closed after 100 minutes, and there was no significant difference from normal gap closure. c, d, Time-lapse images of gap closure when ROCK was inhibited with 25 µM Y-27632. Scale bars are 100 µm.

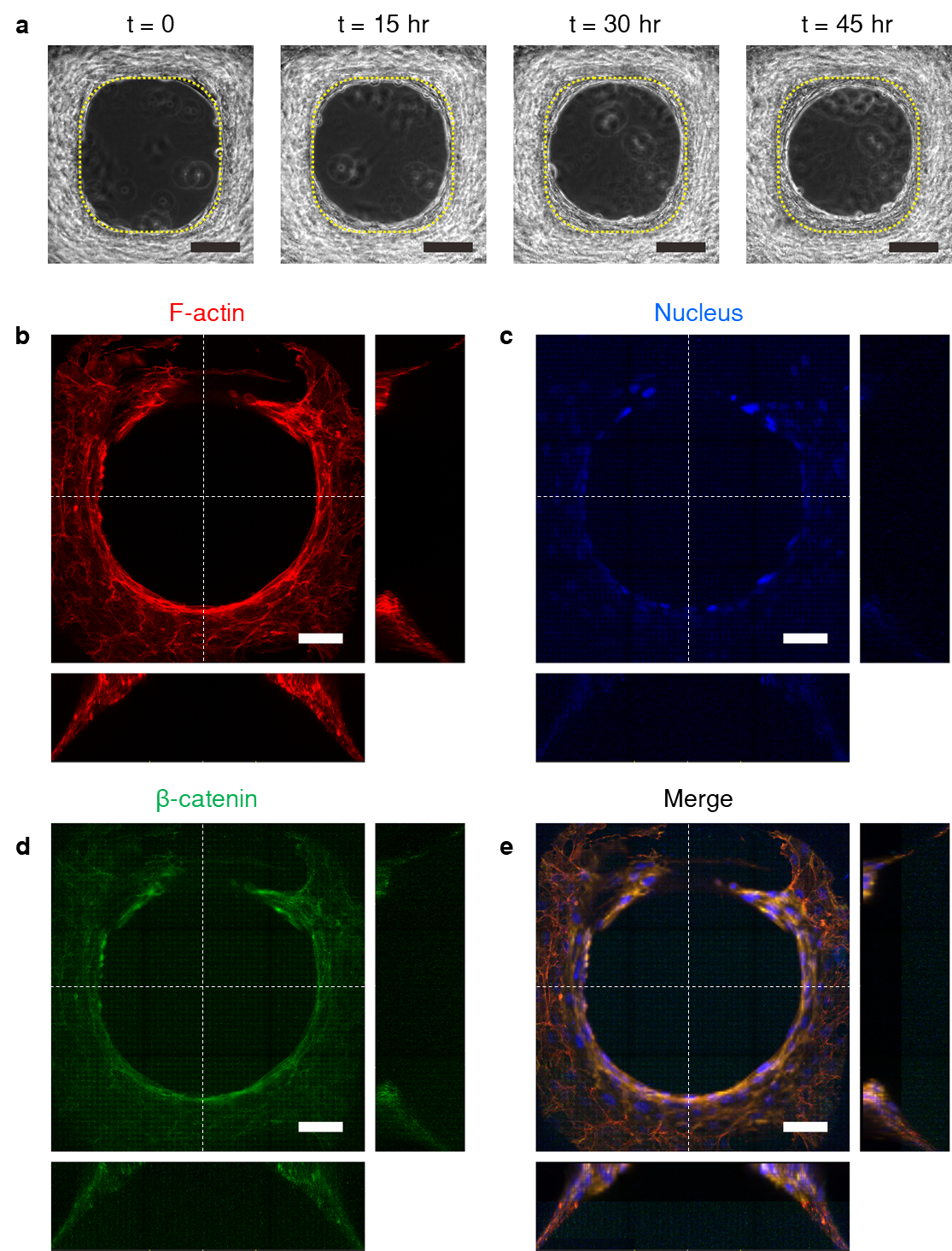

**Supplementary Fig. 7.** ROCK-mediated void closure on platform a, Phase contrast images with different time points; the dashed yellow line is the boundary of the void. b-e, Immunofluorescence staining of F-actin (red) (b), nuclei (blue) (c), and β-catenin (green) (d) of cells when inhibited with 25 µM Y-27632 after cells were partially microvoided. (e) Merged images of F-actin, nuclei and adherens junctions from (b-d). Scale bars are 50 µm.

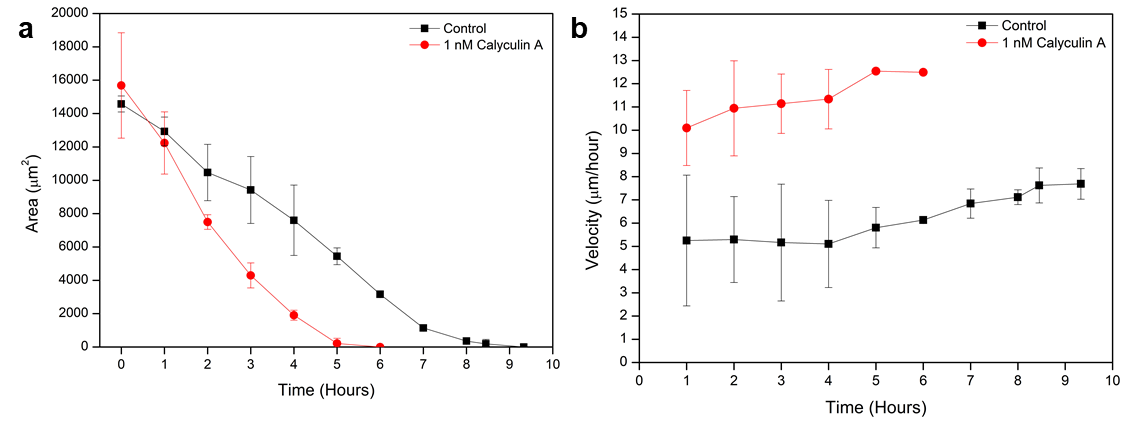

**Supplementary Fig. 8.** Dependence of contractility of myosin II. The closure rate of cells treated with calyculin A on rectangular 3D microvoids was drastically increased. a, The decay of area with time. b, The velocity of cell migration with 1 nM calyculin A treatment.

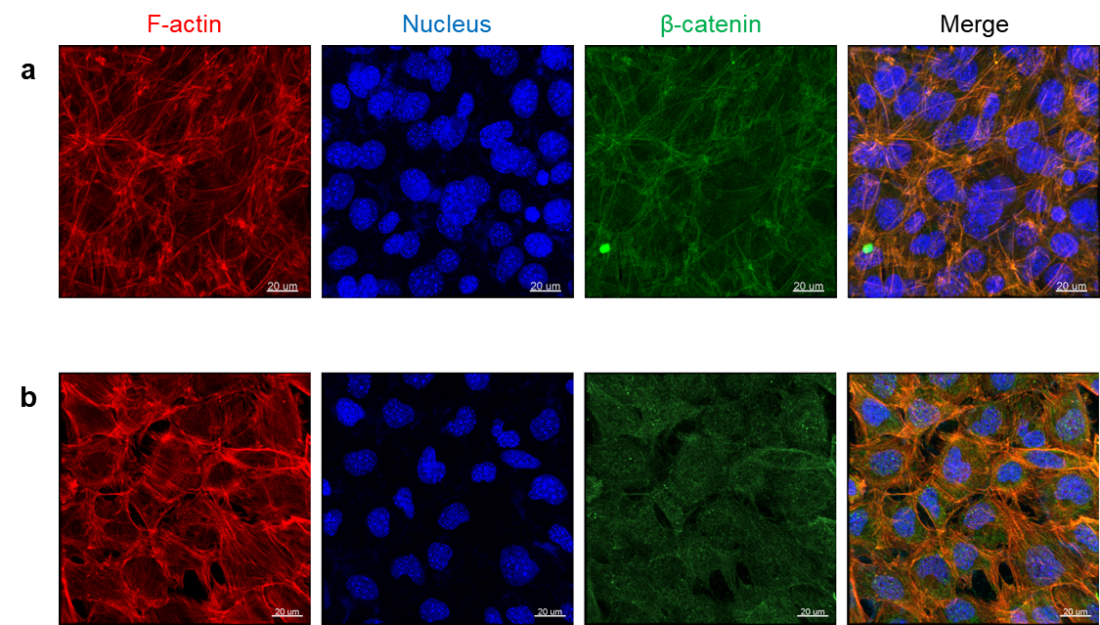

**Supplementary Fig. 9.** Confirmation of the inhibition of adherens junctions on 2D substrates. a, Cells without the inhibition of adherens junctions as a control. b, Adherens junctions were inhibited by using low Ca^2+^ DMEM. The result shows the absence of intercellular adherens junctions between the cells. Scale bars are 20 µm.

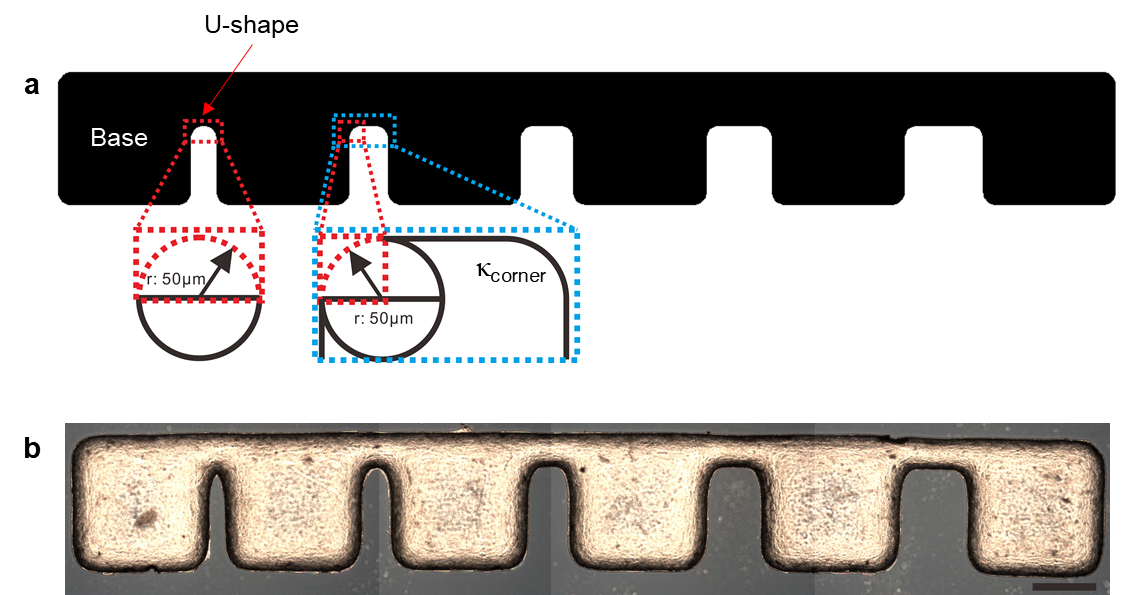

**Supplementary Fig. 10a.** Pattern design of open contour 3D linear microgaps with varying gap sizes (100 - 500 µm diameter). b, A PDMS-based 3D linear microgap taken using an optical microscope. The scale bar is 500 µm.

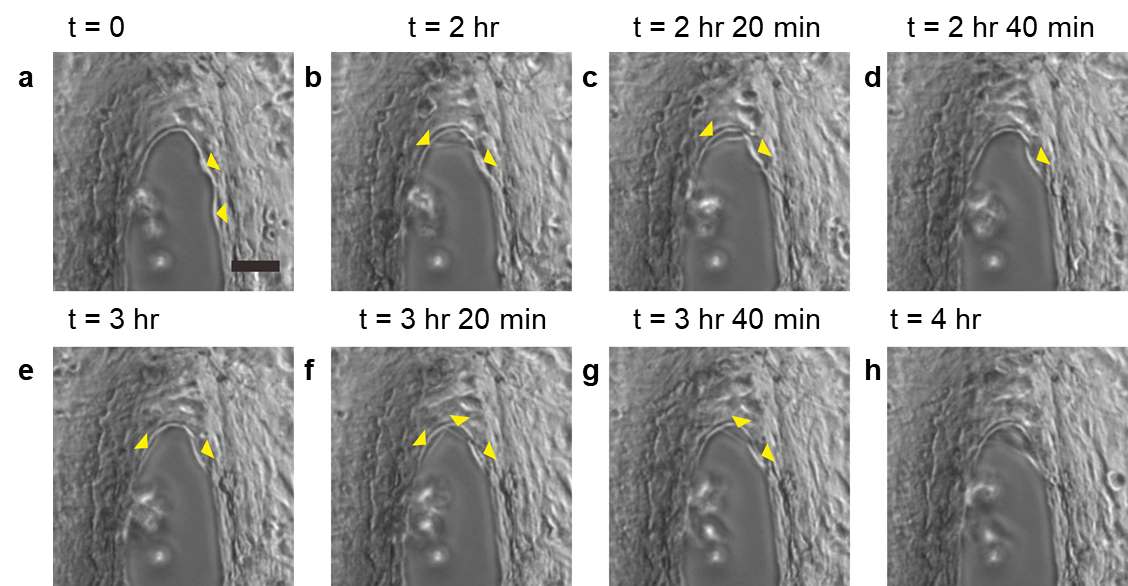

**Supplementary Fig. 11.** Initiation of fibroblast bridge formation in a 100 µm gap shown in a 3D linear microgap from 0 h to ~4 h. a, Two individual fibroblast cells were observed trying to migrate to the gap from the middle (yellow head arrows). b, One of the cells retracted from the middle; however, the other cell approached from the curvature of the U-shaped structure. c, Cells from the middle remained on the curvature of the U-shape, and little retraction was observed from the curvature of the U-shape. d, The retracted fibroblast was completely withdrawn from the curvature of the U-shaped structure, whereas the other fibroblast maintained its location in the middle of the gap. e, Another cell was observed trying to migrate towards the gap from the middle, and f, this cell migrated to the curvature of the U-shape. A third cell approached the two cells from the curvature of the U-shape. g, One cell retracted, leaving two cells on the curvature of the U-shape. h, The cells began building a stable adherens junction and began to migrate towards the gap from the curvature of the U-shape. The scale bar is 50 µm.

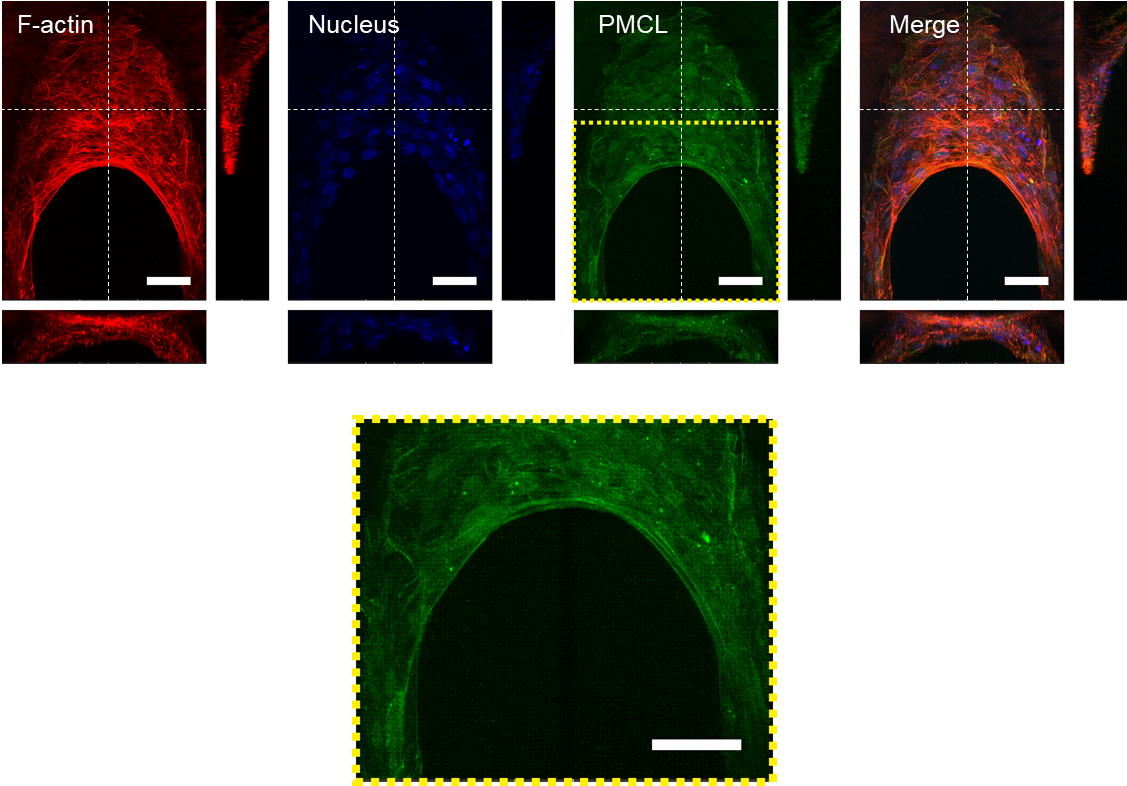

**Supplementary Fig. 12.** Microgap closure mediated by purse string contraction during the zip-up mechanism. The first row of images show 3D reconstructions of immunofluorescence staining for F-actin (red), nuclei (blue) and PMLC (green) along the white dotted lines. The second row shows the magnified image from the highlighted first row stained for PMLC. As shown, the expression of PMLC is well aligned along the periphery at the κ_advancing cells_. Scale bars are 50 µm.

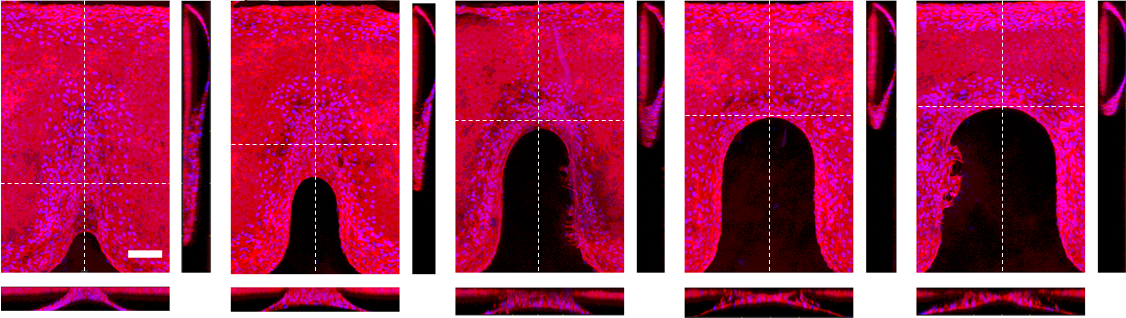

**Supplementary Fig. 13.** Immunofluorescence staining for F-actin (red) and nuclei (blue) after 85 hr of zip-up gap closure in 3D linear gaps of various widths from 100 to 500 µm. The scale bar is 100 µm.

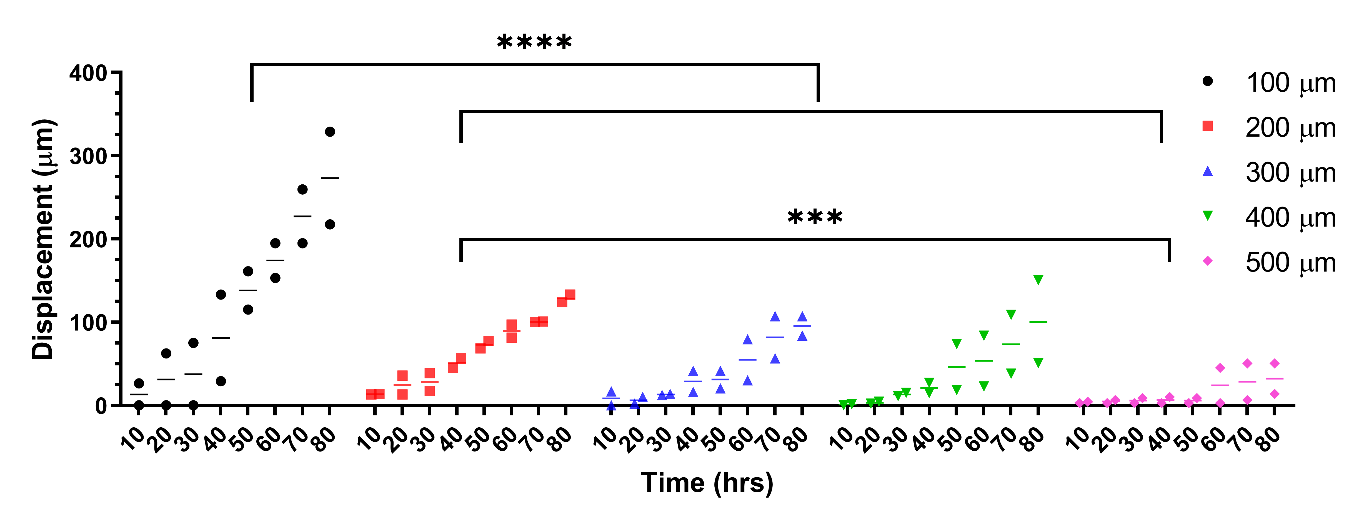

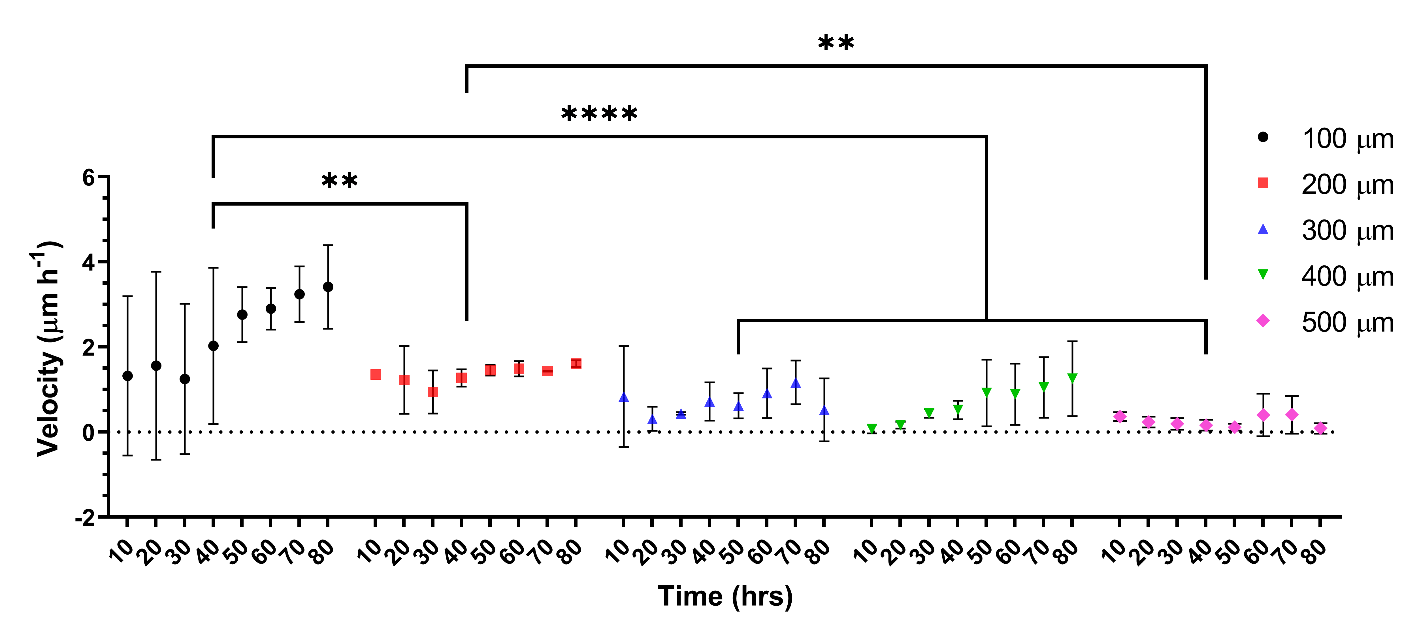

**Supplementary Fig. 14.** Displacement and velocity of zip-up gap closure of gaps of various widths (100-500 µm). The results are the mean ± standard deviation (SD) of triplicate measurements and are representative of three independent experiments. ***** p < 0.0001, *** p < 0.001* and *** p < 0.01* indicate a statistically significant difference.

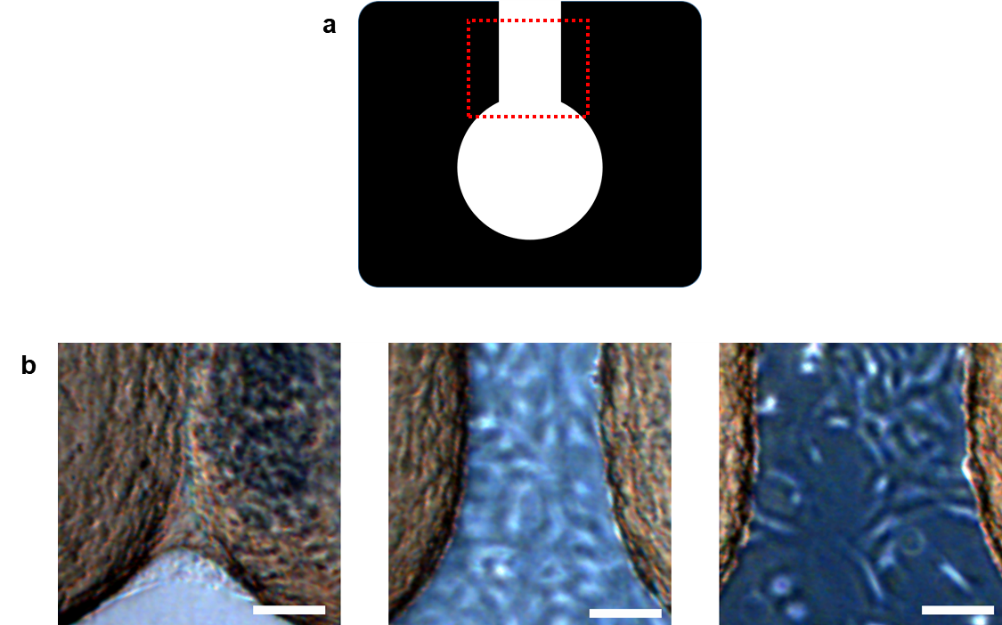

**Supplementary Fig. 15a.** Pattern design of the 3D microgap without a U-shaped structure. b, Optical microscope images of cells at the highlighted area in (a) after 50 hr of cell culture with gap widths of 10, 100 and 200 µm. The results indicate that there is no bridge formation after the U-shaped structure is removed except for the 10 µm gap width. The scale bar is 100 µm.

**Supplementary movies**

**Supplementary Movie 1.** NIH 3T3 mouse fibroblast cells close a 100 µm diameter of 3D microvoid.

**Supplementary Movie 2.** NIH 3T3 mouse fibroblast cells close a 200 µm diameter of 3D microvoid.

**Supplementary Movie 3.** NIH 3T3 mouse fibroblast cells close a 300 µm diameter of 3D microvoid.

**Supplementary Movie 4.** NIH 3T3 cell gap closure is mediated by contractility of myosin II before the initiation of void closure.

**Supplementary Movie 5.** NIH 3T3 cell gap closure is mediated by contractility of myosin II after partial void closure.

**Supplementary Movie 6.** Role of ROCK on a smaller void.

**Supplementary Movie 7.** Role of ROCK on a larger void.

**Supplementary Movie 8.** Adherens junctions of NIH 3T3 cells were inhibited before void closure.

**Supplementary Movie 9.** Adherens junctions of NIH 3T3 cells were inhibited after partial void closure.

**Supplementary Movie 10.** Zip-up gap closure on an open contour 3D linear microgap.

**Supplementary Movie 11.** 3D reconstruction of fibroblast bridge formation from confocal images during the zip-up mechanism.
